# More than *Mycobacterium tuberculosis*: specific site-of-disease microbial communities, functional capacities, and their distinct clinical profiles in tuberculous lymphadenitis

**DOI:** 10.1101/2022.06.16.496073

**Authors:** Georgina Nyawo, Charissa Naidoo, Benjamin Wu, Imran Sulaiman, Jose Clemente, Yonghua Li, Stephanie Minnies, Byron Reeve, Suventha Moodley, Cornelia Rautenbach, Colleen Wright, Shivani Singh, Andrew Whitelaw, Pawel Schubert, Robin Warren, Leopoldo Segal, Grant Theron

## Abstract

**Background:** Lymphadenitis is the most common extrapulmonary tuberculosis (EPTB) manifestation and a major cause of death. The microbiome is important to human health but uninvestigated in EPTB. We profiled the site-of-disease lymph node microbiome in tuberculosis lymphadenitis (TBL).

**Methods:** Fine needle aspiration biopsies (FNABs) were collected from 159 pre-treatment presumptive TBL patients in Cape Town, South Africa. 16S Illumina MiSeq rRNA gene sequencing was done.

**Results:** We analysed 89 definite TBLs (dTBLs) and 61 non-TBLs (nTBLs), which had similar α-but different β-diversities *(p=0.001).* Clustering identified five lymphotypes prior to TB status stratification: *Mycobacterium-, Prevotella*- and *Streptococcus*-dominant lymphotypes were more frequent in dTBLs whereas a *Corynebacterium-dominant*lymphotype and a fifth lymphotype (no dominant taxon) were more frequent in nTBLs. When restricted to dTBLs, clustering identified a *Mycobacterium-dominant* lymphotype with low α-diversity and other non-*Mycobacterium*-dominated lymphotypes (termed *Prevotella-Corynebacterium* and *Prevotella-Streptococcus*). The *Mycobacterium* dTBL lymphotype was associated with HIV-positivity and clinical features characteristic of severe lymphadenitis (e.g., node size). dTBL microbial communities were enriched with potentially proinflammatory microbial short chain fatty acid metabolic pathways (propanoate, butanoate) vs. those in nTBLs. 11% (7/61) of nTBLs had *Mycobacterium* reads.

**Conclusions:** TBL at the site-of-disease is not microbially homogenous and distinct microbial community clusters exist that are associated with different immunomodulatory potentials and clinical characteristics. Non-*Mycobacterium*-dominated dTBL lymphotypes, which contain taxa potentially targeted by TB treatment, represent less severe potentially earlier stage disease. These investigations lay foundations for studying the microbiome’s role in lymphatic TB and the long-term clinical significance of lymphotypes requires prospective evaluation.

## INTRODUCTION

Tuberculosis (TB), which kills 1.5 million people globally each year (including 214 000 people with HIV), causes extrapulmonary tuberculosis (EPTB)^1^. EPTB accounts for ~15% of all TB, and as much as half of all TB in in people living with HIV (PLHIV) in some settings^2^. EPTB is difficult to diagnose^3^ and has high mortality.

TB lymphadenitis (TBL) is the most common EPTB manifestation, accounting for 70% of EPTB and most frequently affects peripheral and cervical lymph nodes^4^. TBL occurs after *Mtb* enters the airways, is taken up by phagocytic cells, and transported to a thoracic lymph node where granulomas may form. These steps are also necessary for priming T-cells to generate adaptive immune responses for microbial killing mediated by cytokines and other effector mechanisms^5 6^.

Lymph nodes have an important role in TB pathogenesis: enlargement has been documented following exposure, even if only a fraction of patients with enlarged nodes develop active disease^7^. Furthermore, *Mtb* DNA is often found in the lymph nodes of exposed yet healthy people. Lymph nodes are therefore hypothesised to serve as a *Mtb* growth and persistence niche^7^ that can spread to bodily sites^8^ (in animals lymph node infection almost always accompanies infection in the lungs^9 10^); suggesting that TB may primarily be a lymphatic rather than pulmonary disease^11^. For example, prior to development of active TB, lymph nodes demonstrate enhanced metabolic activity on PET-CT scans^12^. Together these studies show the lymph nodes have an important role in TB pathogenesis, however, the determinants of why *Mtb* sometimes successfully establishes itself in the lymph nodes and subsequently proliferates, including the potential role of other microbes, is understudied. Key to understanding this is characterizing the local site-of-disease.

The human microbiome influences immune function^13^. Two studies assessed lymph node microbial content^14 15^, both in mesenteric lymph nodes in Crohn’s disease where reduced diversity was observed. The site-of-disease microbiome in TB is underexamined^16^: in bronchoalveolar lavage fluid (BALF), active pulmonary TB was associated with *Mycobacterium* enrichment and *Streptococcus* depletion^17 18^.

The site-of-disease microbiome in TBL (including in HIV-endemic settings where TB is common) remains uncharacterised. Therefore, given the apparent role of the lymph nodes in TB pathogenesis, and the importance of the microbiome as a modulator of immunity, we characterised the site-of-disease lymph microbiome in presumptive TBL patients from a high HIV burden setting before the potentially confounding effects of antibiotic-based TB treatment.

## METHODS

### Patient recruitment and follow-up

Presumptive TBL participants (≥18 years) were recruited from Tygerberg Academic Hospital in Cape Town, SA (25 January 2017-11 December 2018). Participants were programmatically referred for a routine fine needle aspiration biopsy (FNAB) for the investigation of lymphadenopathy as described^19^. Eligible participants were not on TB treatment within six months. Clinical and demographic data were collected by interview and medical record review. Patients with TBL were programmatically diagnosed with TB, initiated on treatment, and study staff assessed treatment response by telephonic follow-up ≥12 weeks. The study had no role in patient management.

### Ethics

Patients provided written informed consent. The study was approved by the Health Research and Ethical Committee of Stellenbosch University (N16/04/050), Tygerberg Hospital (Project ID:4134), and the Western Cape Department of Health (WC_2016RP15_762).

### Specimen collection and processing

For each patient, two background DNA sampling controls were collected in microcentrifuge tubes prior to lymph node aspiration: a skin swab (collected into saline; Ysterplaat Medical Supplies, Cape Town, South Africa) of the site to be punctured, followed by a saline flush of the syringe to be used for aspiration. Aspiration and microbiological procedures are in the Supplement. Aspirated material from the third pass was collected into 500μL sterile saline and stored at −80 °C until batched DNA extraction.

### Routine specimen testing

Patients were categorised based on lymphatic or non-lymphatic mycobacteriological evidence, provided by the government programmatic laboratory (National Health Laboratory Service [NHLS]), and/or clinical decision to start treatment (**Figure 1**) by responsible clinician thereafter. Case definitions are described in the Supplement (**Table S1**).

**Figure 1:**
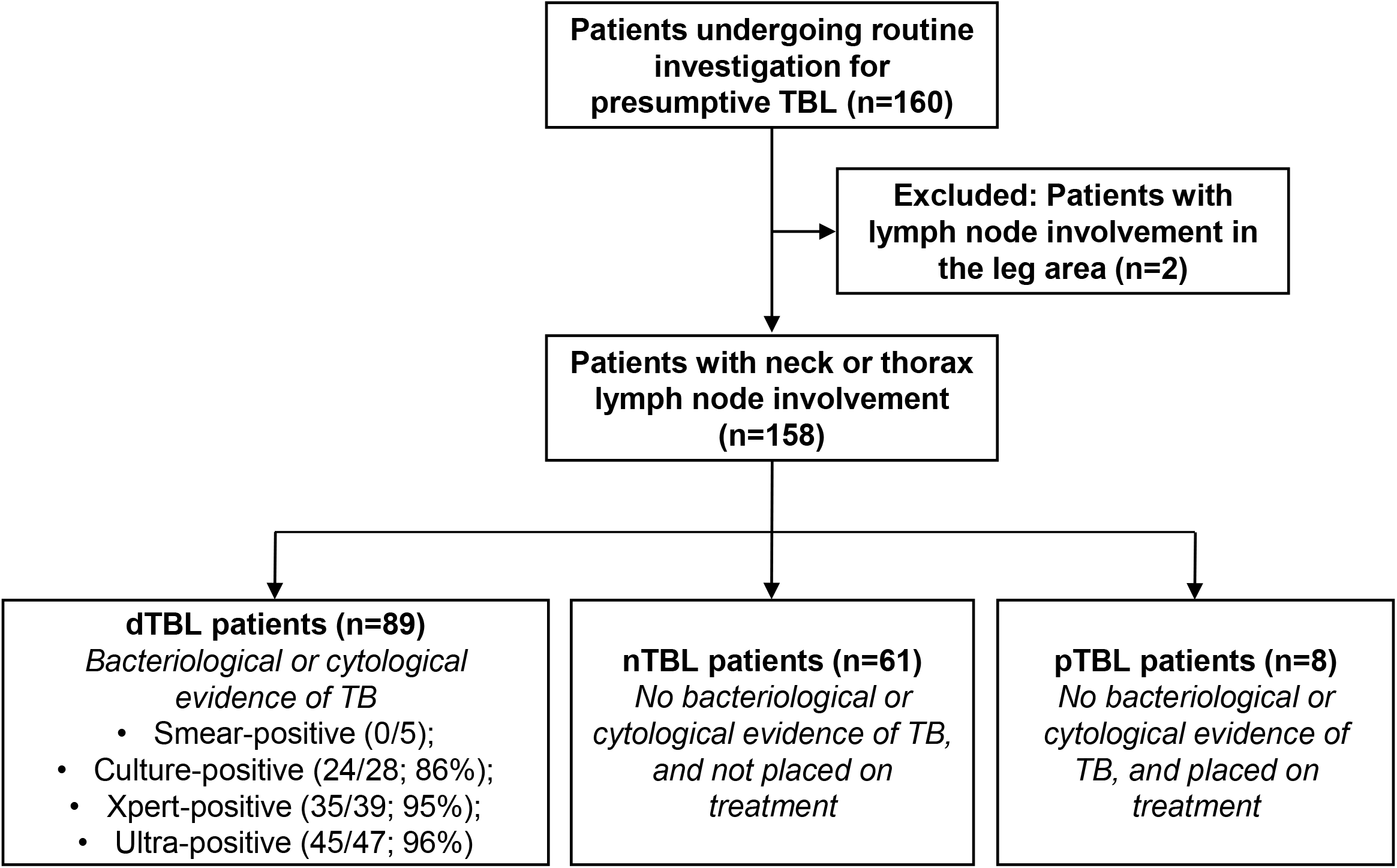
Study flow chart. Fine needle aspirates, skin and saline controls were collected from presumptive TBL patients. dTBLs: definite tuberculous lymphadenitis; nTBLs: non-tuberculous lymphadenitis; pTBLs: probable tuberculous lymphadenitis; Smear: Smear microscopy; MGIT960 Culture: Mycobacteria Growth Indicator Tube 960 liquid culture; Xpert: Xpert MTB/RIF; Ultra: Xpert MTB/RIF Ultra.

### Microbial DNA extraction and sequencing

DNA was extracted from specimens and controls using the PureLink Microbiome DNA Purification Kit (Invitrogen, Carlsbad, USA). The 16S rRNA gene V4 hypervariable region (150 bp read length) was amplified and sequenced (paired-ends) on the Illumina MiSeq platform. Lymph, skin swab, and 1 in 5 saline flushes were extracted and sequenced.

### Microbiome data analysis

16S rRNA gene sequences ([Sequence Read Archive](https://www.ncbi.nlm.nih.gov/sra) PRJNA738676) were processed, denoised and analysed in Quantitative Insights Into Microbial Ecology (QIIME 2, v2020.8)^20^and DADA2^21^ using closed-reference picking by assigning taxonomy at a 97% similarity against representative sequences in Greengenes (v13.8)^22^. QIIME2 outputs (phylogenetic tree, feature table, taxonomy) and metadata imported into R (v3.5.2) and analyses done using *phyloseq*^23^ Shannon’s index was calculated using *vegan*^24^ to measure α-diversity (within-sample diversity). Bray-Curtis distances were used to measure β-diversity (between-sample diversity) and construct Principal Coordinate Analysis (PCoA) plots. Dirichlet-Multinomial Mixtures (DMM) modelling was done to estimate the optimal number of clusters (microbial community states known as lymphotypes) based on compositional similarity^25^.

### Inferred metagenome

Phylogenetic Investigation of Communities by Reconstruction of Unobserved States (PICRUSt) v.2.1.3-b^26^ was used to predict gene family abundance with PICRUSt2 default options (picrust2_pipeline.py). The resulting gene table was mapped against the Kyoto Encyclopaedia of Genes and Genomes (KEGG) database, and pathway abundances were inferred from predicted KEGG ORTHOLOGY (KO) abundances.

### Differential abundance analysis

Differentially abundant taxa and pathways were identified using *DESeq2* (v1.22.2)^27^ with Benjamini-Hochberg multiple testing correction. Feature tables were pruned to taxa with ≥5% relative abundance in 0.5% of samples^28^ Adjusted p-values <0.2 and <0.05 was considered significant for taxa and pathways, respectively (<u>DESq2 tables</u>). PICRUSt *DeSeq2* outputs were used to identify common pathways in L4 vs. each lymphotype (overall patients), and L3 vs. each other lymphotype (dTBLs).

### Statistical analyses

Statistical analysis was done in GraphPad Prism Version 7.00 (GraphPad Software, USA) and R. For α-diversity comparisons, the non-parametric Mann-Whitney test or the Kruskal-Wallis ANOVA with Dunn’s multiple comparisons was used. For paired comparisons, the Wilcoxon signed rank test was used. Permutational multivariate analysis of variance (PERMANOVA) was computed with 999 permutations to test β-diversity differences and *R*^2^ used to measure the proportion variation explained by a variable. For correlation analysis, the non-parametric Spearman ranking and parametric Pearson ranking tests were used. A p-value <0.05 was significant for all comparisons.

## RESULTS

### Cohort characteristics

We had 89 dTBLs, 61 nTBLs (**Figure 1**) and 9 pTBLs (latter subsequently excluded due to small numbers). dTBLs were more likely to have supraclavicular or head lymph node involvement than nTBLs, if HIV-positive were more likely to have a lower CD4 count (**Table 1**) and were more likely to have a FNAB that appeared bloody rather than chylous.

**Table 1:** Demographic and clinical characteristics of patients with presumptive TBL. dTBLs were more likely to have HIV and a lower CD4 count if HIV-positive, supraclavicular lymph node involvement, and a bloody FNAB. Data are n/N (%) or median (IQR).

|  | Patients with presumptive TB (n=158) |  |  | p-value |
| --- | --- | --- | --- | --- |
|  | Total (n=158) | dTBL (n=89) | nTBL (n=61) |  |
| Age, years | 36 (21-44) | 35 (29-40) | 38 (30-49) | 0.053 |
| Female | 85/159 (53) | 48/89 (54) | 35/61 (57) | 0.677 |
| HIV | 77/156 (49) | 49/89 (55) | 23/59 (39) | 0.055 |
| CD4+ | 166 (90-308) | 155 (76-251) | 250 (139-458) | **0.027** |
| CD4+ <200 cells/μl | 47/77 (61) | 32/49 (65) | 11/23 (48) | 0.159 |
| On ART | 38/76 (50) | 21/49 (43) | 14/22 (64) | 0.105 |
| Previous TB | 36/156 (23) | 24/88 (27) | 9/60 (15) | 0.078 |
| Tobacco smoking | 44/157 (28) | 21/89 (24) | 22/60 (37) | 0.084 |
| Antibiotic use within 1 year of recruitment | 41/155 (26) | 22/87 (25) | 16/60 (27) | 0.851 |
| At recruitment | 24/41 (59) | 10/22 (45) | 11/16 (69) | 0.154 |
| Lymph node characteristics: sites |  |  |  |  |
| Neck | 138/158 (87) | 78/89 (88) | 55/61 (90) | 0.632 |
| Deep anterior cervical | 65/138 (47) | 36/78 (46) | 24/55 (47) | 0.774 |
| Deep lateral cervical | 25/138 (18) | 15/78 (19) | 10/55 (18) | 0.879 |
| Superficial | 15/138 (11) | 6/78 (8) | 9/55 (16) | 0.120 |
| Supraclavicular | 19/138 (14) | 16/78 (21) | 3/55 (5) | **0.015** |
| Head | 13/138 (10) | 4/78 (5) | 9/55 (16) | **0.032** |
| Thorax | 20/158 (13) | 11/89 (12) | 6/61 (10) | 0.632 |
| Axillary (vs. breast) | 16/21 (81) | 9/11 (82) | 3/5 (60) | 0.350 |
| Lymph node characteristics: size, cm<sup>2</sup> | 4 (2-9) | 4 (2-9) | 4 (4-9) | 0.150 |
| Specimen appearance |  |  |  |  |
| Bloody (vs. chylous) | 130/158 (82) | 66/89 (72) | 57/61 (93) | **0.003** |

Bolded items indicate that p values are significant at p<0.05
\*8 Probable TBLs (pTBLs) excluded from table
Abbreviations: dTBLs: definite tuberculous lymphadenitis; nTBLs: non-tuberculous lymphadenitis; pTBLs: probable-tuberculous; ART: Antiretroviral therapy.
<sup>†</sup>Missing data: HIV (n=2); Specimen appearance (n=3)

### Lymph microbiome is distinct from background sampling controls

Lymph fluid had similar α-diversity to background controls (skin, saline) but different β-diversity resulting from an enrichment of *Mycobacterium* **(Figure S1A-D)**, suggesting environmental contamination unlikely.

### Mycobacterium enrichment in dTBLs drives differences with nTBLs

α-Diversity was similar in dTBLs and nTBLs (**Figure 2A**) and, in β-diversity analyses, *Mycobacterium* was the most discriminatory taxon (**Figure 2B-C**) appearing at several fold higher frequencies than in nTBLs (**Figure 2D**). Bray distances within nTBLs were greater than within dTBLs (**Figure 2E**), thus dTBLs were more like each other than nTBLs to each other (likely reflecting the mixture of different disease pathologies in the nTBLs and relative homogeneity of dTBLs). *Mycobacterium* reads were present in 64% (57/89) of dTBLs and 11% (7/61; *p<0.0001)* of nTBLs (**Figure S2**) and, when sequences underwent BLAST, all reads matched with *Mtb*.

**Figure 2:**
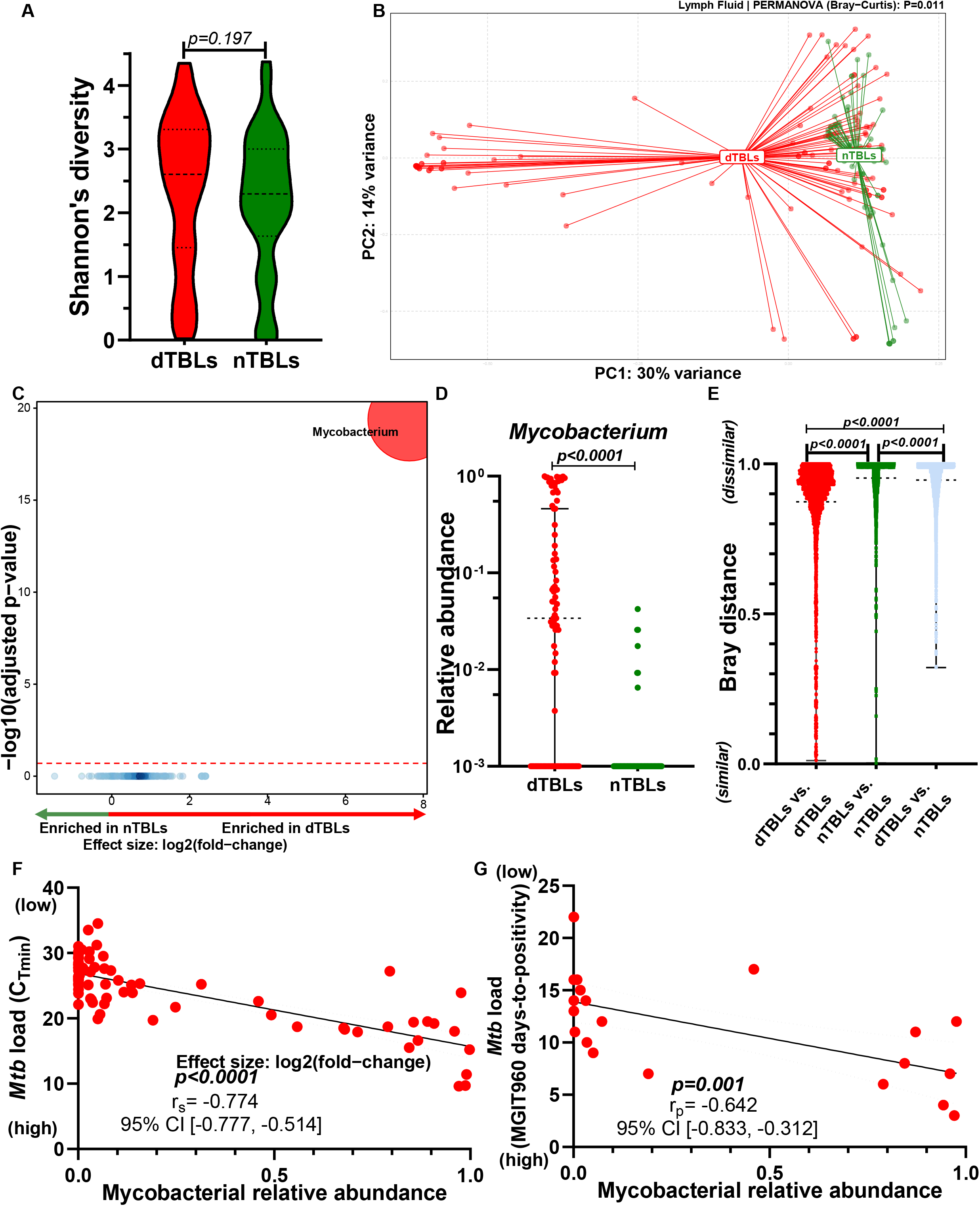
dTBLs have a distinct microbiome to nTBLs with *Mycobacterium* enrichment. **(A)** Although α-diversity was similar, **(B)** β-diversity differed. *Mycobacterium* was enriched in dTBLs compared to nTBLs based on **(C)**differential abundance testing and **(D)**relative abundance. Discriminatory taxa appear above the threshold (red dotted line, FDR=0.2). **(E)** dTBLs were more compositionally similar to each other than nTBLs. Mycobacterial reads positively correlated with *Mtb* load: **(F)** Xpert and Ultra and **(G)** culture (days-to-positivity). dTBLs: definite tuberculous lymphadenitis; nTBLs: non-tuberculous lymphadenitis; Xpert: Xpert MTB/RIF; Ultra: Xpert MTB/RIF Ultra; MGIT960: Mycobacteria Growth Indicator Tube 960 liquid culture; r_s_: Spearman correlation coefficient; r_p_: Pearson correlation coefficient.

### Correlation between 16S rRNA gene sequencing and TB diagnostic test results

As expected, there was a higher relative abundance of *Mycobacterium* reads in dTBLs [median 0.034 (IQR 0.001-0.460) vs. 0.001 (0.001-0.001), *p<0.0001;* **Figure 2D**]. *Mycobacterium* relative abundance in dTBLs showed a positive correlation with bacillary load (based on Xpert and Ultra cycle threshold values; r_s_=-0.774, 95% CI [−0.777, −0.514], *p<0.0001;***Figure 2F**), and culture days-to-positivity; r_p_=-0.642, [−0.833, −0.31.], *p=0.001;***Figure 2G**). Furthermore, a non-significant trend towards a positive correlation between lymph node size and mycobacterial load (relative abundance, Xpert/Ultra C_Tmin_ values) was observed (**Figure S3A-B**).

### Differences by HIV status

Overall: α-diversity did not differ by HIV status (**Figure 3A**) and although β-diversity did (**Figure 3B**) no differentially enriched taxa were found (not shown), however, the relative abundance of *Mycobacterium* was higher in PLHIV (**Figure S4A**).

**Figure 3:**
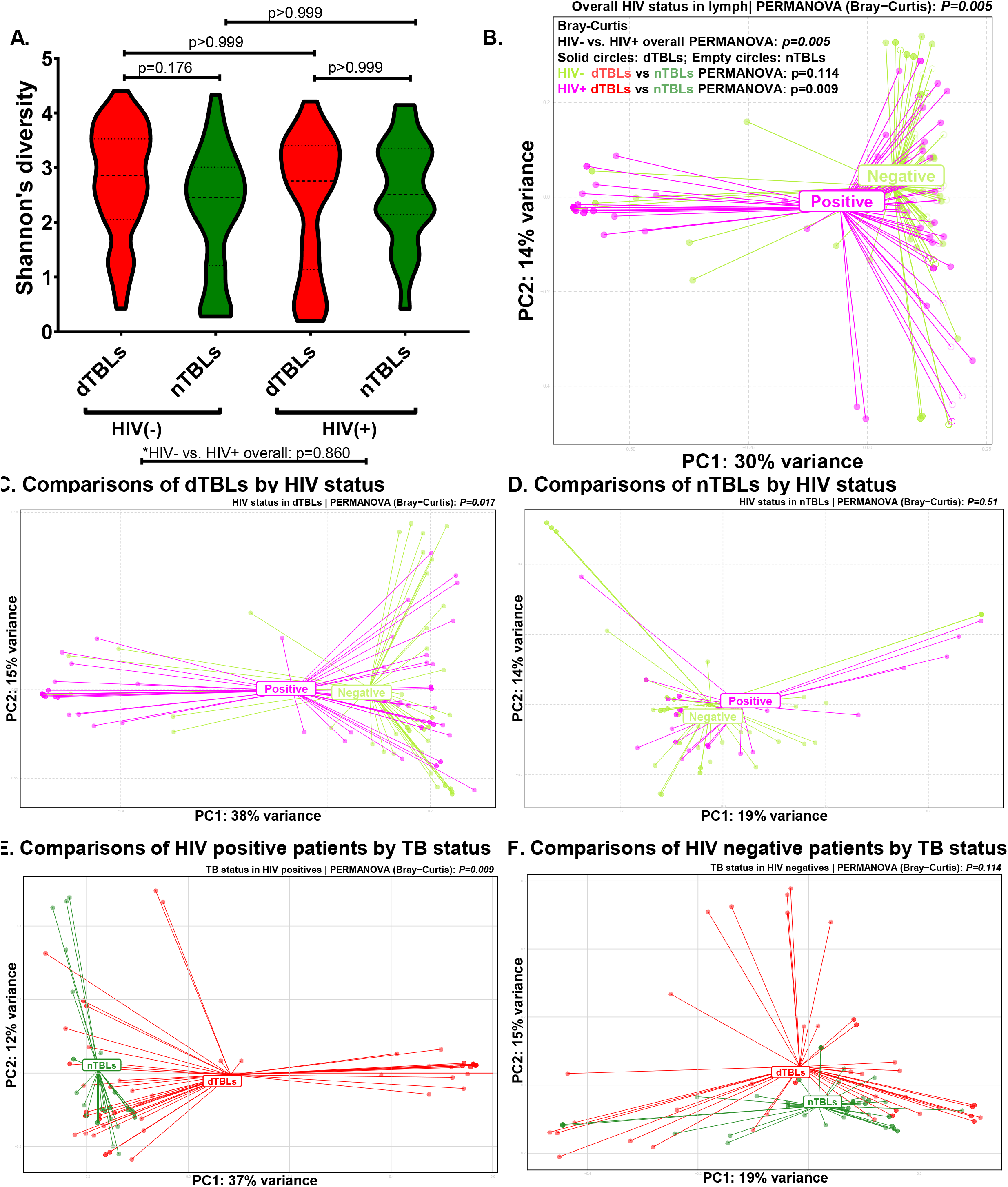
Microbiome differences in HIV-positive dTBLs versus nTBLs but not in HIV-negative dTBLs versus nTBLs. **(A)** α-Diversity did not differ by HIV or TBL statuses, **(B)** however, β-diversity differed between HIV-positives and -negatives overall (shaded circles are dTBLs, empty circles are nTBLs). β-diversity differed by HIV status in **(C)** dTBLs but not in **(D)** nTBLs. β-diversity differed by TBL status in **(E)**HIV-positives and **(F)** HIV-negatives.

Comparisons within dTBLs or nTBLs by HIV status: There were 55% (49/89) and 39% (23/59) HIV-positive dTBLs and nTBLs, respectively. Within dTBLs or nTBLs, α-diversities did not differ by HIV status (**Figure 3A**) and β-diversity differed within dTBLs by HIV status *(p=0.017,***Figure 3C**) but not within nTBLs. HIV-positive dTBLs had higher *Mycobacterium* relative abundance than HIV-negative dTBLs **(Figure S4A)**.

Comparisons within HIV-positives or -negatives by TB status: In people with the same HIV status, α-diversity did not differ by TB status (**Figure 3A**) and β-diversity only differed between dTBLs vs. nTBLs in HIV-positives *(p=0.009,***Figure 3E**) where dTBLs were enriched in *Mycobacterium* (**Figure S4B**). In HIV-negatives, there were no differences between dTBLs and nTBLs (**Figure 3F**).

### Lymphotype identification and their associations with clinical characteristics

Overall: Five lymphotypes with differing α- and β-diversities were identified (**Figure 4A-C, Table S3**). L1 had no dominant taxa (**Figure 4D**), whilst L4 was *Mycobacterium-dominated* and had the least α-diversity, and L2, L3 and L5 were *Corynebacterium-, Prevotella-* and *Streptococcus-*dominated, respectively. While no taxa were differentially abundant in L1 vs. other lymphotypes (**Figure S5A-C**), L2, L3, and L5 were enriched relative to L4 in *Corynebacterium, Prevotella,* and *Streptococcus,* respectively (**Figure 4E-G**). The proportions of dTBLs in L1, L2, L3, L4, and L5 were 35% (17/48), 63% (28/44), 57% (12/21), 100% (21/21), and 69% (11/16), respectively. The patients in these lymphotypes are associated with distinct clinical characteristics. The majority of nTBLs occurred in highly diverse lymphotypes with a heterogenous mixtures of taxa; likely reflecting the spectrum of pathologies in people with TBL ruled out. L1 was associated with characteristics indicative of less severe lymphadenitis. Compared separately to L2, L4, and L5, L1s were less likely to have dTBL. Furthermore, L1s were less likely to be HIV-positive vs. L4s but, L1 PLHIVs had lower CD4 counts vs. L2 and L3 PLHIVs. In contrast, L4 was associated with characteristics resembling more severe lymphadenitis. L4 was more likely to contain dTBL patients than each other lymphotype. Furthermore, compared to L2s, L4s were more likely to have a bigger lymph node, chylous FNABs and, of PLHIV, a smaller proportion on ART. Compared to L3s, L4s were more likely to have previous TB and HIV, and those with HIV were more likely to have lower CD4 counts. Compared to L5s, L3s with HIV had lower CD4 counts. Therefore, in summary, L1 appears to be associated with less severe forms of lymphadenitis, whereas L4 was associated more severe forms (**Table S4**).

**Figure 4:**
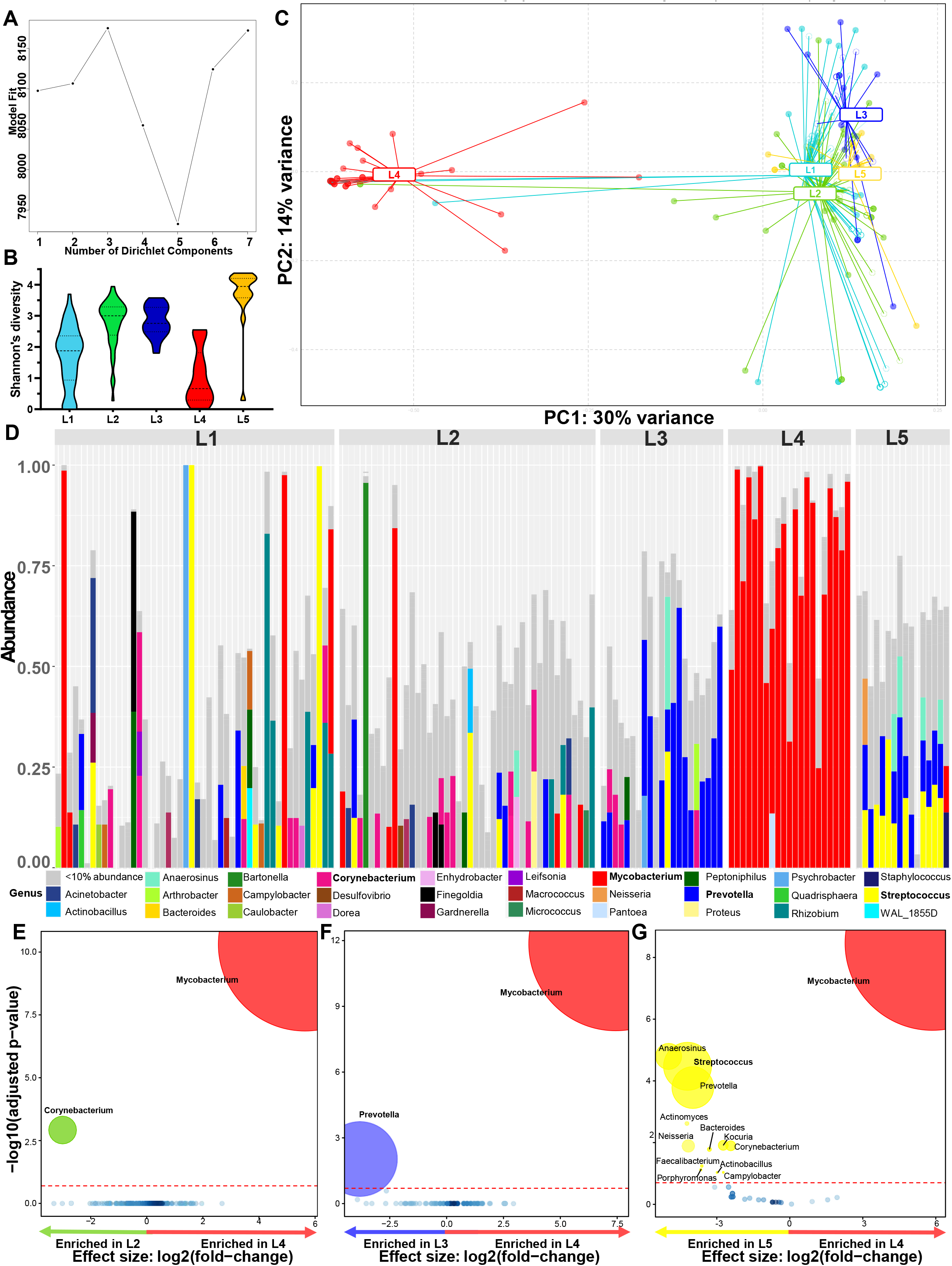
Five lymphotypes observed in presumptive TBL. **(A)** LaPlace approximation identified five clusters. **(B)** L5 had the highest α-diversity. **(C)**β-diversity differed between each lymphotype (shaded circles dTBLs, empty circles nTBLs). **(D)** Stacked bar plots showing L1 with a heterogenous mixture of genera, L2 dominated by *Corynebacterium,* L3 dominated by *Prevotella*, L4 dominated by *Mycobacterium*, and L4 dominated by *Streptococcus*. Bolded taxa represent dominating taxa. **(E)** *Corynebacterium* was enriched in L2; **(F)** *Prevotella* enriched in L3, **(G)** *Mycobacterium* enriched in L4, and *Streptococcus* enriched in L5. Significantly more discriminatory taxa (bolded) appear closer to the left or right, and higher above the threshold (red dotted line, FDR=0.2) as significance increases. Relative taxa abundance is indicated by circle size. dTBLs: definite tuberculous lymphadenitis; nTBLs: non-tuberculous lymphadenitis; L: lymphotype.

Within patients of the same TB status: Within dTBLs, three lymphotypes with differing β-diversities were identified (**Figure 5A-B**). L1 was abundant in *Prevotella* and *Corynebacterium,* L2 in *Prevotella* and *Streptococcus,* and L3 in *Mycobacterium* (**Figure 5C**) and these taxa were differentially abundant (**Figure 5D-F**). These lymphotypes were termed *Prevotella-Corynebacterium*, *Prevotella-Streptococcus* and *Mycobacterium,* respectively. L3s were more likely to be HIV-positive, with larger lymph nodes, compared to L1s. In addition, L3s were more likely to have larger lymph nodes than L2s. Lastly, L2s are more likely to be female than L1s (**Table S5**). Together, these differences suggest L3 is associated with more severe TBL than other lymphotypes. Within nTBLs, no lymphotypes were identified (**Figure S6**).

**Figure 5:**
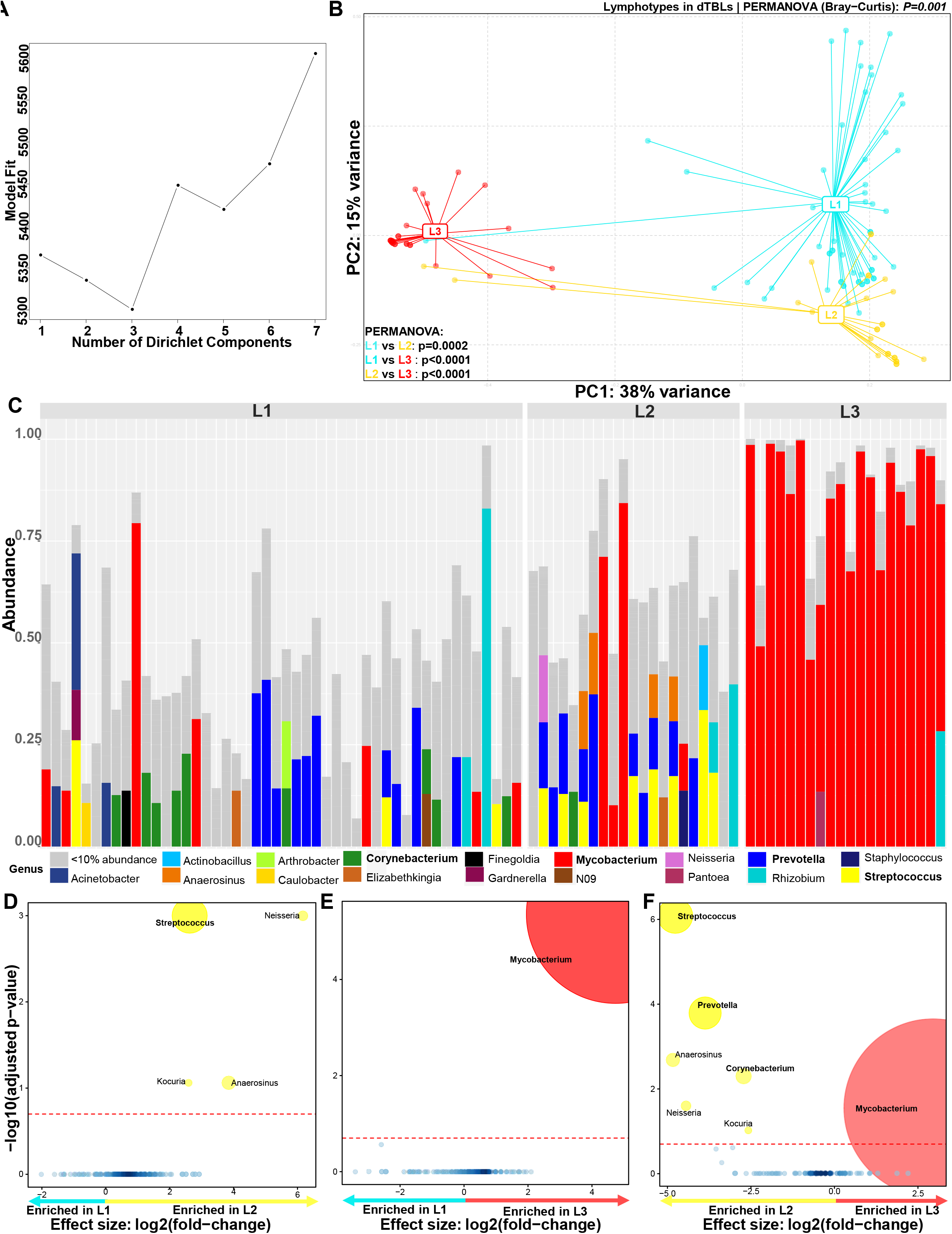
Three lymphotypes identified in dTBLs. **(A)** Best model fit based on LaPlace approximation identified three clusters within dTBLs. **(B)** β-diversity differed between lymphotypes. **(D)** Stacked bar plots showing L1 comprised of *Mycobacterium* and accompanying heterogenous taxa, L2 dominated by *Prevotella* and *Streptococus*, and L3 dominated by *Mycobacterium.* Bolded taxa represent dominating taxa. **(D)**No taxa were enriched in L1, **(E)** L2 was enriched in *Streptococcus,* **(F)** and *Mycobaterium* was enriched in L3. Significantly more discriminatory taxa (bolded) appear closer to the left or right, and higher above the threshold (red dotted line, FDR=0·2) as significance increases. Relative taxa abundance is indicated by circle size. dTBLs: definite tuberculous lymphadenitis; L: lymphotype.

### Predictive metagenome profiling shows increased short chain fatty acid metabolism

dTBLs vs. nTBLs: 139 inferred microbial metabolic pathways were differentially enriched (75 in dTBLs, 64 in nTBLs). In dTBLs, “fatty acid metabolism”, “benzoate degradation”, “propanoate metabolism” and “butanoate metabolism” were enriched, suggesting increased SCFA production (**Figure 6**).

**Figure 6:**
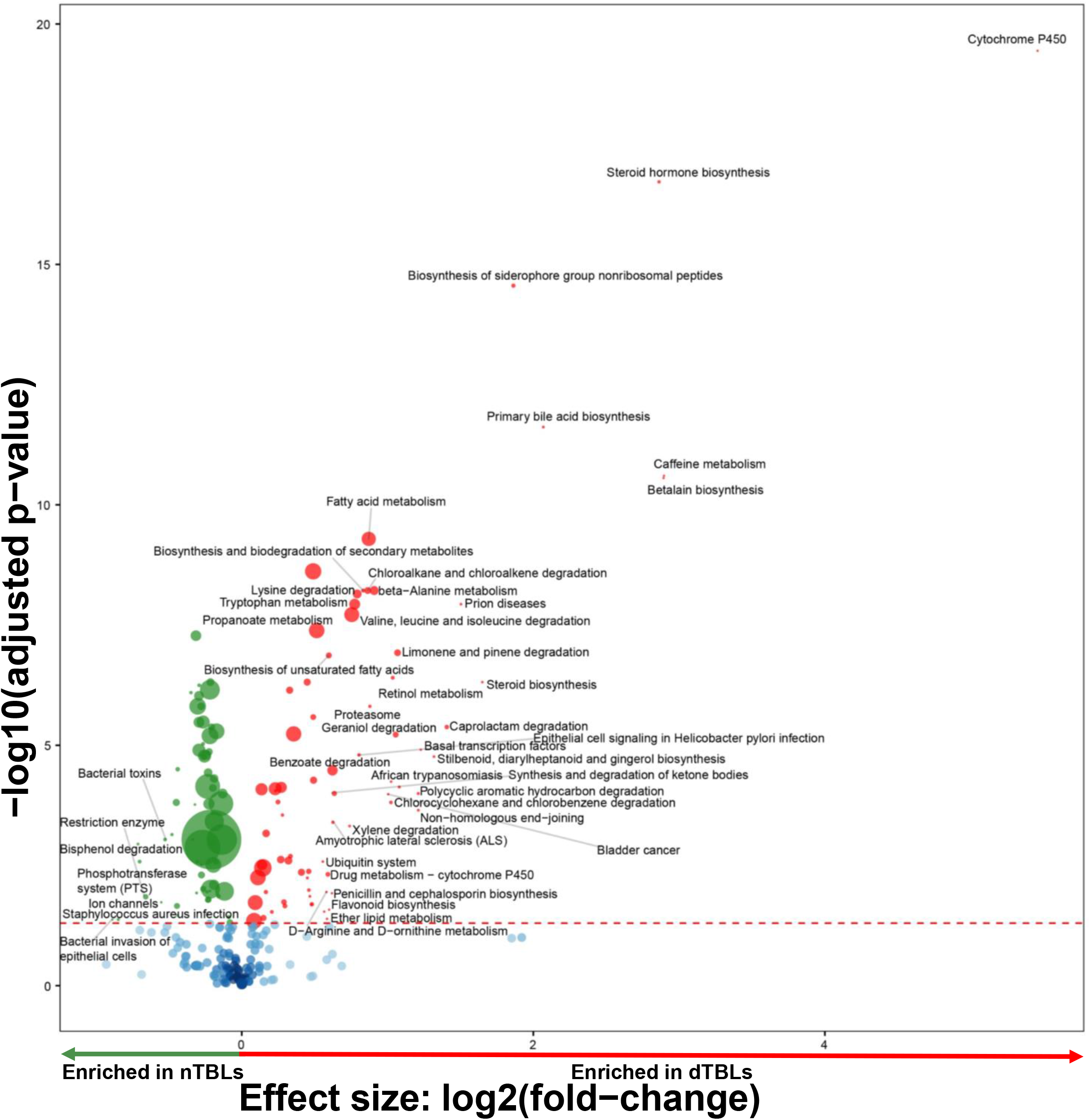
Enriched microbial capacity for SCFA pathways in dTBLs versus nTBLs. Volcano plot depicting differentially abundant microbial pathways in dTBLs vs. nTBLs inferred by PICRUSt2. Key pathways of interest are bolded including aminobenzoate degradation, benzoate degradation, and propanoate degradation. Significantly more discriminatory pathways appear closer to the left or right, and higher above the threshold (red dotted line, FDR=0.05) as significance increases. Relative gene abundance is indicated by circle size. SCFA: short chain fatty acids; dTBLs: definite tuberculous lymphadenitis; nTBLs: non-tuberculous lymphadenitis.

HIV-positive vs. negatives: The above SCFA-related pathways were enriched in HIV-positive vs. -negative patients overall and, within dTBLs, in HIV-positives vs. -negatives (**Figure 7A-B**). Within nTBLs, HIV-positives were enriched in the “cell cycle – *Caulobacter”*, “bacterial secretion system” and “oxidative phosphorylation” vs. -negatives (**Figure S7**).

**Figure 7:**
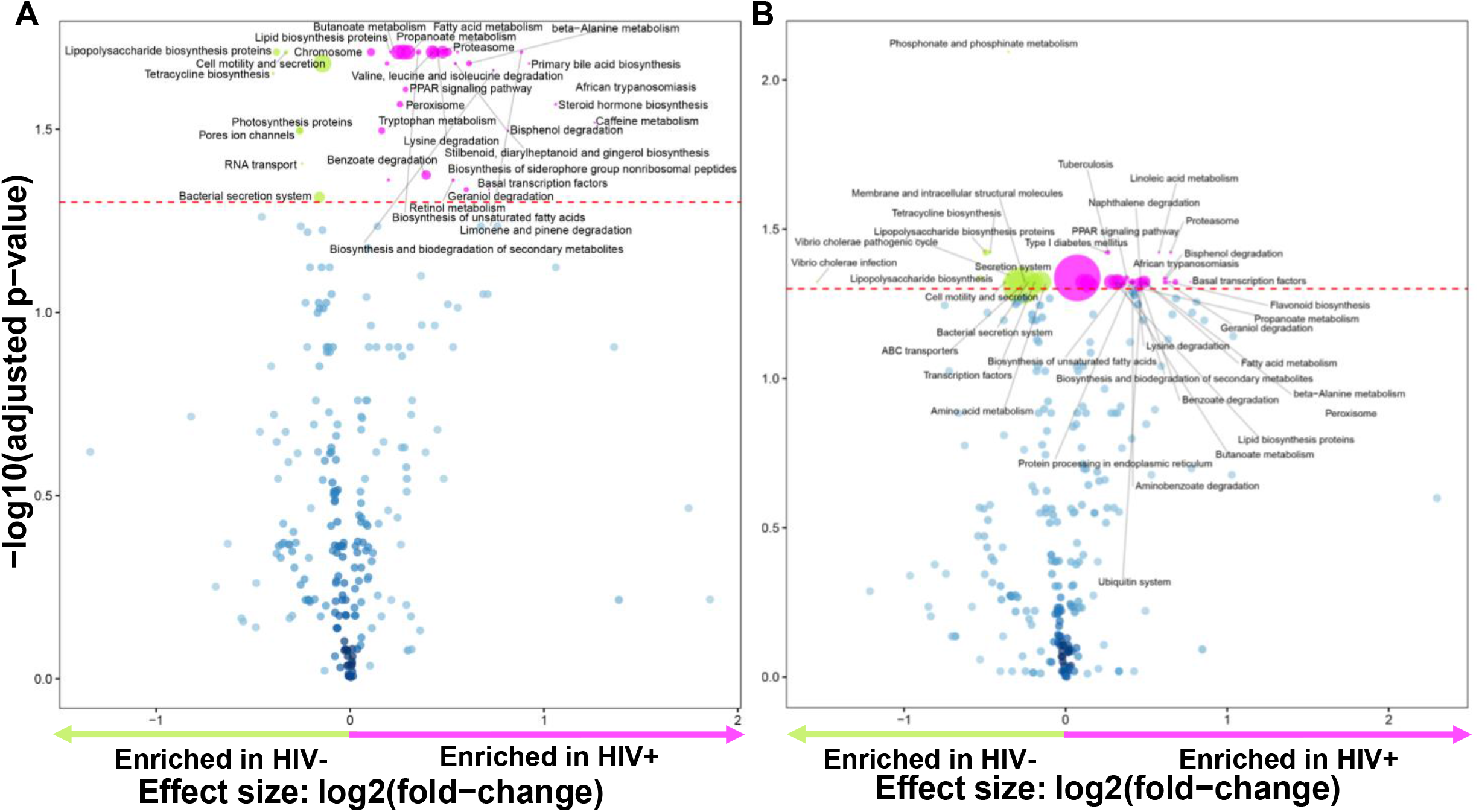
Predicted metagenome function reveals increased capacity for SCFA production in HIV-positive versus HIV-negative patients overall, and in dTBLs. Volcano plot depicting functional pathways differing between **(A)** HIV-positive and HIV-negative patients with presumptive TBL, and **(B)** in dTBLs. Key pathways of interest include butanoate metabolism, propanoate metabolism and benzoate degradation. Significantly more discriminatory pathways appear closer to the left or right, and higher above the threshold (red dotted line, FDR=0.05). Relative gene abundance is indicated by circle size. SCFA: short chain fatty acids.

In different lymphotypes: When comparing lymphotypes’ inferred pathways in all patients (overall including dTBLs and nTBLs), a similar core of pathways was enriched in lymphotype 4. These included the “propanoate metabolism”, “tuberculosis”, “lipid biosynthesis”, “butanoate metabolism”, “fatty acid metabolism” and “PPAR signalling pathway” (most-to-least enriched) (**Figure 8A-B)**. In contrast, vs. lymphotype 4, lymphotype 1 was enriched in “epithelial cell signalling in *Helicobacter pylori* infection”, lymphotype 2 was enriched in “carbohydrate digestion and absorption”, lymphotype 3 was enriched in “dioxin degradation”, and lymphotype 5 was enriched in “carbohydrate digestion and absorption” (**Figure S8A-H)**. When comparing the three dTBL lymphotypes, *Mycobacterium*-dominated lymphotype 3 was, compared to each other dTBL lymphotypes, enriched in the similar core pathways seen for the *Mycobacterium-*dominated lymphotype 4 overall in all patients (**Figure 8C; Figure S9**).

**Figure 8:**
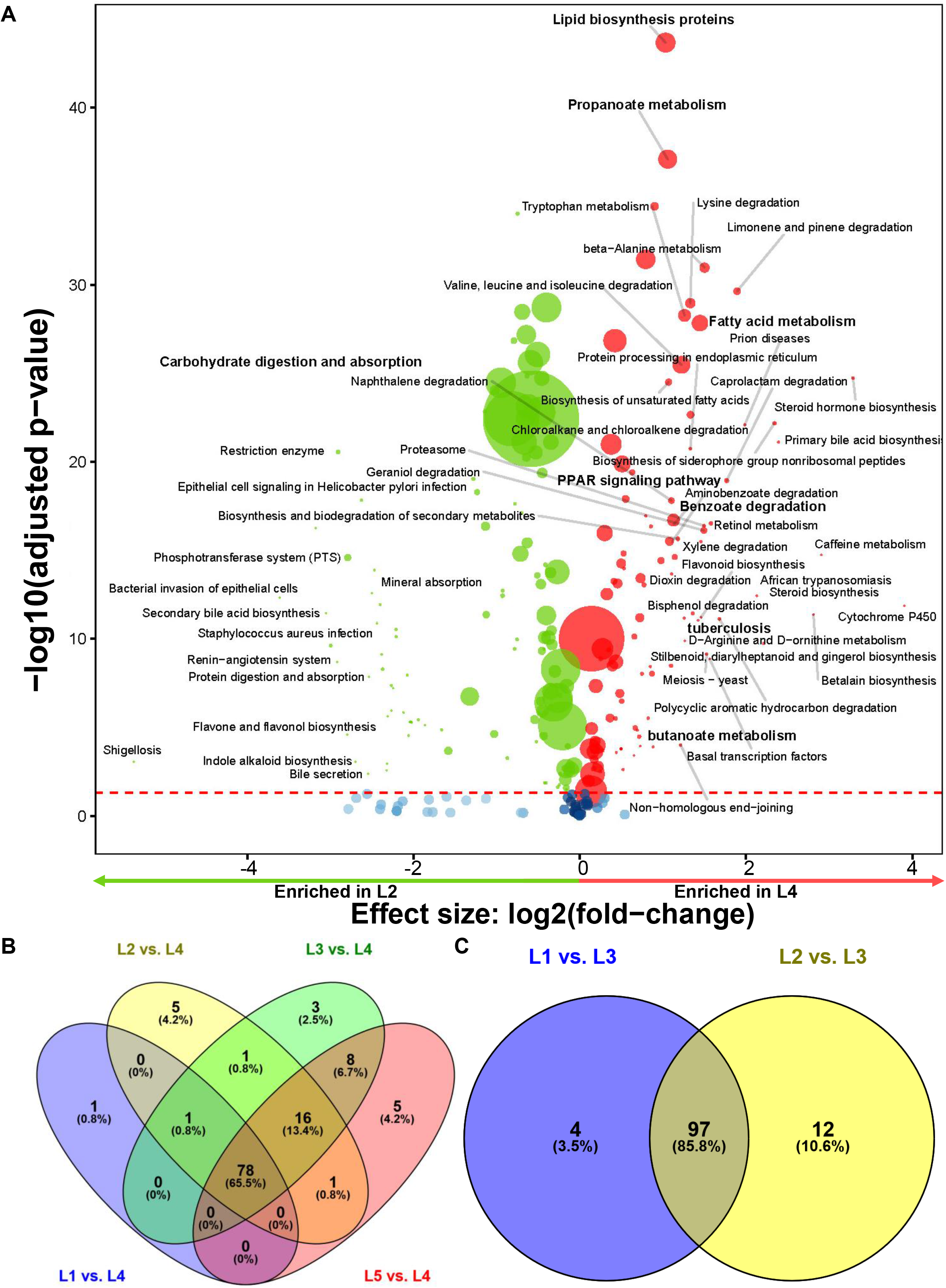
Differential microbial pathways between lymphotypes showing similar core pathways enriched in the *Mycobacterium-dominated* lymphotype.. (A) Volcano plot showing differentially abundant microbial pathways inferred by PICRUSt2 in lymphotype 2 vs. lymphotype 4 representing pathways enriched in lymphotype 4 compared to every other lymphotype in all patients (overall including dTBLs and nTBLs). Significantly more discriminatory pathways appear closer to the left or right, and higher above the threshold (red dotted line, FDR=0.05) as significance increases. Relative gene abundance is indicated by circle size. **(B)** 65.5% of all inferred pathways enriched L4 compared to each other overall lymphotypes were common, whilst **(C)** 85.8% were common in L3 compared to each other dTBL lymphotypes. Differentially enriched pathways common in all comparisons with the *Mycobacterium* dominant lymphotype included pathways involving lipid biosynthesis, fatty acids, and SCFA metabolism i.e. lipid biosynthesis proteins, propanoate metabolism, benzoate degradation, and valine, leucine and isoleucine degradation. SCFA: short chain fatty acids; dTBLs: definite tuberculous lymphadenitis; nTBLs: non-tuberculous lymphadenitis; L: lymphotype.

## DISCUSSION

We characterised the local microbial environment in patients with lymphadenitis undergoing investigation for TB in a HIV-endemic setting. Our key findings are: 1) lymphatic microbial communities in dTBLs clustered into three distinct “lymphotypes” we termed *“Prevotella-Corynebacterium”, “Prevotella-Streptococcus”,* and *“Mycobacterium”,* 2) the *Mycobacterium* dTBL lymphotype was associated with HIV-positivity and other clinical features characteristic of severe lymphadenitis, and 3) dTBLs relative to nTBLs were functionally enriched in fatty acid-, amino acid-, and SCFA-related microbial metabolic pathways with known immunomodulatory effects (the *Mycobacterium* lymphotype was most enriched in these pathways than other dTBL lymphotypes). Finally, 4) dTBLs without *Mycobacterium* reads and nTBLs with *Mycobacterium* reads were identified. These data show TBL at the site-of-disease is not microbially homogenous and that distinct clusters of microbial communities exist associated with different clinical characteristics. The long-term significance and importance of these lymphotypes requires prospective evaluation.

We identified three lymphotypes within dTBLs termed “*Prevotella-Corynebacterium”, “Prevotella-Streptococcus”*, and “*Mycobacterium”*, distinguished by different relative abundances of these taxa (*Prevotella* co-occurred in the first two lymphotypes). These individual taxa are enriched in respiratory secretions from pulmonary TB cases^29 30^. Furthermore, within dTBLs, *Streptococcus* is associated with low BMI and extent of lung damage^30^. *Prevotella* in bronchoalveolar lavage fluid also positively correlates with SCFA concentrations and independently predicts incident TB in people without co-prevalent TB^31^. Compared to the other dTBL lymphotypes, “*Mycobacterium”* was associated with severe disease and most frequently occurred in PLHIV, agreeing with diagnostics studies that show stronger baseline mycobacterial PCR test readouts predict long term clinical outcomes in pulmonary^32^ and extrapulmonary TB^33^. Together, these data show distinct lymphotypes are associated with different clinical characteristics and suggests that patients with the most severe *Mycobacterium-dominated* lymphotype may initially progress through different site-of-disease microbial states characterised by *Corynebacterium-, Streptococcus-* and/or *Prevotella*-domination. Studies with longitudinal follow-up and repeat sampling are required to examine whether these lymphotypes have potential for clinical staging.

Importantly, *Corynebacterium* and *Streptococcus* often dominated in dTBL patients. Members of both taxa are causative agents of lymphadenitis and, even though these patients have TB lymphadenitis confirmed via conventional diagnostics, *Corynebacterium* and *Streptococcus* may therefore co-contribute to pathology and symptoms^34–36^. Coincidently, these taxa fall within the anti-microbial spectrum of first-line TB treatment^16^, meaning that this regimen may, in part, cure lymphadenitis by killing *Corynebacterium* and *Streptococcus* in addition to *Mycobacterium.*

Microbial pathways predicted to be most enriched in dTBLs involved fatty acid, amino acid, and SCFAs (benzoate, propanoate) metabolism; all of which are associated with pulmonary TB disease compared to sick patients without TB^37 38^. SCFAs in particular suppress immune pathways involved in IFN-γ and IL-17A production and, *ex vivo,* limit macrophage-mediated kill of *Mtb.* SCFA concentrations hence predict incident TB in patients^31^. Our research therefore suggests that the inflammation associated with lymphadenopathy is in part caused by the presence of microbes including but limited to *Mycobacterium* that are able to produce SCFAs that interfere with these immunological pathways; revealing potentially new therapeutic targets to reduce lymphadenopathy.

We detected *Mycobacterium* in nTBLs. These reads could be from previous TB exposure or disease. *Mtb* DNA has been found in the lymph nodes of healthy individuals and primates exposed to TB, where the sites are hypothesised to serve as a *Mtb* growth and persistence niche^7^. dTBLs without *Mycobacterium* reads were also documented, however, 16S rRNA sequencing has well known sub-optimal sensitivity for *Mycobacterium*, in part due to low 16s RNA gene copy number^39^.

Our study has strengths and limitations. Patients were sampled once, as close as possible to treatment initiation; animal models might permit repeat invasive sampling especially if treatment is withheld. We did not co-analyse host immune signatures but plan to do so. The programmatic context enabled large numbers of patients to be recruited, however, detailed long-term follow-up, which could include imaging of lymph nodes and more detailed measurements of differential responses to treatment, was not possible.

In conclusion, we show dTBL patients have a distinct microbiome at the site of disease, characterized by three lymphotypes (*Mycobacterium, Prevotella-Corynebacterium, Prevotella-Streptococcus)*. This dysbiosis of the lymphatic microbiome likely contributes to pathophysiology, including inflammatory state and clinical severity, which itself may reflect the chronicity of TB disease. TB lymphadenitis does therefore not appear to be a microbially homogenesis disease, and this reveals potentially new diagnosis, therapeutic, and prognostic targets.

## ACKNOWLEDGMENTS

The authors thank study participants and Tygerberg Hospital FNA clinic staff especially Sr Cupido. Additionally, we thank CLIME research group staff, especially Sr Ruth Wilson and Roxanne Higgit. GRN acknowledges funding from L’Oréal-UNESCO For Women in Science Sub-Saharan Africa Young Talents Award, and the International Rising Talents Award. The content is the solely the responsibility of the authors and does not necessarily represent the official views of the funders. Computations were performed using facilities provided by the University of Cape Town’s ICTS High Performance Computing team: <u>hpc.uct.ac.za</u>

## CONTRIBUTORSHIP

GN, CCN and GT contributed to conceptualisation and design of the study and supervised the study, funding acquisition, data collection, and wrote the manuscript. GRN, CCN, IS, BGW, IS, JCC, LNS, and GT contributed to data and statistical analysis, and figures. All authors contributed to interpretation of data and editing of the manuscript. As the study guarantor, GT is responsible for the overall content of this manuscript.

## COMPETING INTRESTS

None

## DATA AVAILABILITY

Data are available upon reasonable request De-identified patient data, the study protocol, informed consent, and datasets generated in this study may be requested from the corresponding author.

