## Supplement for "More than *Mycobacterium tuberculosis*: specific site-of-disease microbial communities, functional capacities, and their distinct clinical profiles in tuberculous lymphadenitis"

**Supplementary Material**

### Methods

#### *FNAB collection and TB microbiology*

Needle passes were done on the largest (using surface area recorded in $cm^2$) distinct node with a 23-gauge needle and a 10 mL syringe as described (1). The first two passes were used to prepare standard microscope slides for cytological examination using Rapidiff and Papanicolaou staining. A flush of the needle was collected in 1.5 mL of TB transport medium (2) media and sent to the National Health Laboratory Services (NHLS) microbiology laboratory for Xpert MTB/RIF (Xpert) or Xpert MTB/RIF Ultra (Ultra), Mycobacteria Growth Indicator Tube 960 liquid culture (MGIT960; BD), and acid-fast bacilli (AFB) staining.

#### *Definitions*

We used a reference standard to designate patients as definite-TBLs (dTBLs), probable-TBLs (pTBLs), or non-TBLs (nTBLs) as previously described (1). dTBLs had at least one *Mtb* complex-positive specimen by acid-fast bacilli (AFB) staining microscopy, Xpert MTB/RIF (Xpert) and/or Xpert MTB/RIF Ultra (Ultra), or Mycobacteria Growth Indicator Tube (MGIT) 960 liquid culture (culture). pTBLs did not meet dTBL criteria but commenced treatment empirically. nTBLs had no microbiological TB, were not placed on treatment, and/or had an alternative diagnosis.

#### *Clustering*

We then evaluated for presence of distinct groups of samples based on identification of distinct microbial communities in lymph nodes which we called lymphotypes. Dirichlet multinomial mixture modelling (DMM) was performed using the R package *DirichletMultinomial* to establish clustering within groups (3). Using genus tables, the number of clusters was determined by selecting the number of Dirichlet components that reduced the Laplace approximation of the model (3) (i.e. lower values indicate better fits). Clustering profiles indicate unique groupings, interpreted as “lymphotypes”.

### Results

*Environmental and background controls*

It is important to evaluate possible sources of microbial DNA contamination in low biomass samples such lymph fluid. Pairwise comparisons of α-diversity were similar between saline and skin, and lymph fluid and saline (**Figure S1A**). β-diversity was different between the three fluid types (*p=0.001*; **Figure S1B**), with lymph enriched in the respiratory pathogen *Mycobacterium* (**Figure S1C**) vs. skin, and vs. saline (**Figure S1D**). Skin was enriched with *Psychrobacter* and *Corynebacterium* vs. lymph and saline, respectively (**Figures S1C and S1E**), whilst no taxa were enriched in saline (**Figures S1D-E**).

*α- and β-diversities according to demographic, clinical, and microbiological characteristics* ***(Table S2)***

Overall: Females had a higher α-diversity than males (*p=0.016*), patients who used antibiotics within a year had a lower α-diversity than those who did not (*p=0.003*), and patients with smaller lymph nodes had a higher α-diversity than those with larger nodes (*p=0.001*). β–diversity was different in patients with antibiotic use at recruitment versus none (*p=0.032*) and antibiotic use within one year versus later use (*p=0.020*). Furthermore, within PLHIV, β–diversity differed by ART status (*p=0.042*) and CD4 count stratum (*p=0.038*).

dTBLs: α-diversity was decreased with antibiotic use at recruitment (*p=0.025*) and within one year (*p=0.007*) as well as in larger nodes lymph node size (*p=0.034*). β-diversity also differed by antibiotics usage (concurrent and within one year) and CD4 count stratum in PLHIV (*p=0.034*).

nTBLs: α-diversity was less in males than females (*p=0.003*) and in smokers than non-smokers (*p=0.002*). β-diversity was only associated with specimen appearance (*p=0.047*). No

**Table S1: Reference standard definition used in the study.** Due a small number of pTBLs, they were excluded from analyses.

| | dTBLs | nTBLs | pTBLs |
|---|---|---|---|
| | **Site-of-disease fluid** | | |
| Xpert | ✓ | ✗ | ✗ |
| Ultra | ✓ | ✗ | ✗ |
| MGIT960 Culture | ✓ | ✗ | ✗ |
| Smear microscopy | ✓ | ✗ | ✗ |
| Cytology | ✓ | ✗ | ✗ |
| | **Non-site-of-disease fluid** | | |
| Smear microscopy | ✓ | ✗ | ✗ |
| Xpert | ✓ | ✗ | ✗ |
| Ultra | ✓ | ✗ | ✗ |
| MGIT960 | ✓ | ✗ | ✗ |
| | **Treatment information** | | |
| TB treatment initiated | ✗ | ✗ | ✓ |
| Response to treatment self-reported by patient | ✗ | ✗ | ✓ |

Abbreviations: dTBLs: definite tuberculous lymphadenitis; nTBLs: non-tuberculous lymphadenitis; pTBLs: probable-tuberculous lymphadenitis; Xpert: Xpert MTB/RIF; Ultra: Xpert MTB/RIF Ultra; MGIT960 Culture: Mycobacteria Growth Indicator Tube 960 liquid culture.

**Table S2: α- and β-diversities in presumptive TBL patients when patients with different demographic and clinical characteristics were compared.** Several characteristics, described in the Supplementary Results text, were associated with differing diversities.

| **Characteristics** | **Overall (n=150)** | | | **dTBLs (n=89)** | | | **nTBLs (n=61)** | | |
|---|---|---|---|---|---|---|---|---|---|
| | α-diversity *p-value* (Shannon's Index) | β-diversity | | α-diversity *p-value* (Shannon's Index | β-diversity | | α-diversity *p-value* (Shannon's Index | β-diversity | |
| | | *p-value* (PERMANOVA) | $R^2$ *value* | | *p-value* (PERMANOVA) | $R^2$ *value* | | *p-value* (PERMANOVA) | $R^2$ *value* |
| dTBL | 0.110 | **0.001** | 0.037 | - | - | - | - | - | - |
| Sex | **0.016** | 0.121 | 0.010 | 0.406 | 0.616 | 0.035 | **0.003** | **0.012** | 0.008 |
| HIV | 0.860 | **0.004** | 0.023 | 0.179 | **0.008** | 0.043 | 0.312 | 0.731 | 0.432 |
| *CD4+ <200 cells/µl* | 0.459 | **0.038** | 0.032 | 0.053 | **0.034** | 0.055 | 0.140 | 0.455 | 0.045 |
| *On ART* | 0.662 | **0.042** | 0.030 | 0.267 | 0.344 | 0.022 | 0.306 | 0.267 | 0.055 |
| Previous TB | 0.337 | 0.072 | 0.012 | 0.426 | 0.141 | 0.018 | 0.501 | 0.603 | 0.015 |
| Tobacco smoking | 0.084 | 0.189 | 0.009 | 0.636 | 0.658 | 0.008 | **0.002** | 0.276 | 0.020 |
| Antibiotic use within 1 year of recruitment | **0.042** | **0.020** | 0.015 | **0.025** | **0.012** | 0.036 | 0.547 | 0.212 | 0.022 |
| Antibiotic use at recruitment | **0.003** | **0.032** | 0.061 | **0.007** | **0.025** | 0.141 | 0.062 | 0.064 | 0.115 |
| Site (neck vs. thorax) | 0.220 | 0.134 | 0.010 | 0.128 | 0.142 | 0.018 | 0.809 | 0.830 | 0.011 |
| Specimen appearance (bloody vs. chylous) | 0.213 | 0.068 | 0.012 | 0.771 | 0.198 | 0.016 | **0.020** | **0.047** | 0.778 |
| Lymph node characteristics: size, $cm^2$ | **0.011** | 0.128 | 0.012 | **0.034** | 0.065 | 0.265 | 0.197 | 0.612 | 0.017 |

*$R^2$ provides the proportion of variation explained (e.g., a factor that has a $R^2 = 0.037$, explains 3.7% of the variation in community composition) by β-diversity.

Abbreviations: TB: tuberculosis; TBL: tuberculous lymphadenitis; ART: antiretroviral therapy; dTBLs: definite tuberculous lymphadenitis; nTBLs: non-tuberculous lymphadenitis

**Table S3: Adjusted p-values for α-diversity comparisons between lymphotypes (all patients) measured by Shannon's diversity index.**

| Comparison | Lymphotype with highest α-diversity | Adjusted *p-value* |
|---|---|---|
| **Lymphotype comparisons in all patients** | | |
| L1 vs. L2 | L2 | **<0.0001** |
| L1 vs. L3 | L3 | **0.0012** |
| L1 vs. L4 | L1 | >0.9999 |
| L1 vs. L5 | L5 | **<0.0001** |
| L2 vs. L3 | L2 | >0.9999 |
| L2 vs. L4 | L2 | **<0.0001** |
| L2 vs. L5 | L5 | **0.0329** |
| L3 vs. L4 | L3 | **<0.0001** |
| L3 vs. L5 | L5 | 0.1088 |
| L4 vs. L5 | L5 | **<0.0001** |
| **Lymphotype comparisons in all dTBLs** | | |
| L1 vs. L2 | L2 | **0.001** |
| L1 vs. L3 | L3 | **<0.0001** |
| L2 vs. L3 | L3 | **0.001** |

Definition of abbreviations: L: Lymphotype.

**Table S4: Demographic, clinical, and microbiological differences in each lymphotype (overall in all patients) showing L1 is likely associated with less severe forms of lymphadenitis whereas L4 is associated with more severe forms.** Amongst other differences, L1s were less likely to have dTBL than L2s, L4s, and L5s. Furthermore, L1s were less likely to be HIV-positive vs. L4s. L1 PLHIV had lower CD4 counts vs. L2 and L3 PLHIVs. In contrast, L4s were more likely to be dTBLs than other lymphotypes. Furthermore, compared to L2, L4s had bigger lymph nodes and were more likely to have chylous FNABs and a smaller proportion of PLHIVs on ART. Compared to L3, L4s were more likely to have previous TB and HIV, and L3 PLHIVs were more likely to have lower CD4 counts. Compared to L5, L3 PLHIV had lower CD4 counts.

| **Characteristic¶** | **Total (n=150)** | **L1 (n=48) (No dominant taxa)** | **L2 (n=44) *Corynebacterium*)** | **L3 (n=21) (*Prevotella*)** | **L4 (n=21) (*Mycobacterium*)** | **L5 (n=16) (*Streptococcus*)** | ***p-value* (L1 vs. L2)** | ***p-value* (L1 vs. L3)** | ***p-value* (L1 vs. L4)** | ***p-value* (L1 vs. L5)** | ***p-value* (L2 vs. L3)** | ***p-value* (L2 vs. L4)** | ***p-value* (L2 vs. L5)** | ***p-value* (L3 vs. L4)** | ***p-value* (L3 vs. L5)** | ***p-value* (L4 vs. L5)** |
|---|---|---|---|---|---|---|---|---|---|---|---|---|---|---|---|---|
| Age, years | 36 (30-45) | 35 (29-47) | 37 (32-47) | 31 (28-46) | 37 (34-43) | 36 (28-45) | >0.999 | >0.999 | >0.999 | >0.999 | >0.999 | >0.999 | >0.999 | >0.999 | >0.999 | >0.999 |
| dTBLs | 89/150 (59) | 17/48 (35) | 28/44 (64) | 12/21 (57) | 21/21 (100) | 11/16 (69) | **0.007** | 0.093 | **<0.001** | **0.020** | 0.615 | **0.001** | 0.713 | **0.001** | 0.471 | **0.006** |
| Female | 83/150 (55) | 25/48 (52) | 26/44 (59) | 8/21 (38) | 12/21 (57) | 12/16 (75) | 0.499 | 0.284 | 0.698 | 0.108 | 0.113 | 0.882 | 0.258 | 0.217 | **0.026** | 0.260 |
| HIV | 72/148 (49) | 20/48 (42) | 24/42 (57) | 6/21 (29) | 15/21 (71) | 7/16 (44) | 0.143 | 0.302 | **0.023** | 0.884 | **0.032** | 0.271 | 0.361 | **0.006** | 0.338 | 0.089 |
| *CD4+* | 166 (90-308) | 35 (29-48) | 171 (86-332) | 255 (154-387) | 83 (17-163) | 136 (54-334) | **<0.0001** | **0.001** | 0.733 | 0.103 | >0.999 | 0.620 | >0.999 | 0.223 | >0.999 | >0.999 |
| *CD4+ <200 cells/µl* | 43/72 (60) | 10/20 (50) | 14/24 (58) | 2/6 (33) | 12/15 (80) | 5/15 (33) | 0.580 | 0.473 | 0.069 | 0.324 | 0.272 | 0.163 | 0.129 | **0.040** | >0.999 | **0.010** |
| *On ART* | 35/71 (49) | 9/20 (45) | 16/24 (67) | 3/6 (50) | 5/15 (33) | 2/6 (33) | 0.149 | 0.829 | 0.486 | 0.612 | 0.449 | **0.042** | 0.136 | 0.477 | 0.558 | >0.999 |
| Previous TB | 33/148 (22) | 10/48 (21) | 9/42 (21) | 2/21 (10) | 9/21 (43) | 3/16 (19) | 0.945 | 0.254 | 0.060 | 0.858 | 0.241 | 0.076 | 0.822 | **0.014** | 0.416 | 0.121 |
| Tobacco smoking | 43/149 (29) | 18/48 (38) | 12/44 (27) | 8/21 (38) | 3/21 (14) | 2/15 (13) | 0.296 | 0.936 | 0.054 | 0.062 | 0.377 | 0.245 | 0.232 | 0.079 | 0.082 | 0.875 |
| Antibiotic use within 1 year of recruitment | 38/147 (26) | 11/47 (23) | 9/9 (100) | 4/20 (20) | 9/20 (45) | 5/16 (31) | **<0.001** | 0.760 | 0.077 | 0.533 | **<0.001** | **0.005** | **0.001** | 0.091 | 0.439 | 0.400 |
| *At recruitment* | 21/38 (55) | 8/11 (73) | 5/9 (56) | 1/4 (25) | 6/9 (67) | 1/5 (20) | 0.423 | 0.095 | 0.769 | **0.049** | 0.308 | 0.629 | 0.198 | 0.164 | 0.858 | 0.094 |
| Lymph node characteristics: sites | | | | | | | | | | | | | | | | |
| Neck | 133/150 (89) | 46/48 (96) | 37/44 (84) | 18/21 (86) | 20/21 (95) | 12/16 (75) | 0.058 | 0.136 | 0.911 | **0.013** | 0.865 | 0.201 | 0.421 | 0.293 | 0.410 | 0.074 |
| *Deep anterior cervical* | 60/133 (45) | 19/46 (41) | 16/37 (43) | 8/18 (44) | 13/20 (65) | 4/12 (33) | 0.859 | 0.819 | 0.077 | 0.615 | 0.933 | 0.117 | 0.544 | 0.203 | 0.543 | 0.082 |
| *Deep lateral cervical* | 25/133 (19) | 13/46 (28) | 8/37 (22) | 2/18 (11) | 2/20 (10) | 0/12 (0) | 0.489 | 0.145 | 0.104 | **0.037** | 0.343 | 0.271 | 0.078 | 0.911 | 0.232 | 0.258 |
| *Superficial* | 15/133 (11) | 8/46 (17) | 2/37 (5) | 3/18 (17) | 2/20 (10) | 0/12 (0) | 0.095 | 0.945 | 0.442 | 0.120 | 0.173 | 0.517 | 0.411 | 0.544 | 0.136 | 0.258 |
| *Supraclavicular* | 20/133 (15) | 2/46 (4) | 7/37 (19) | 3/18 (17) | 3/20 (15) | 5/12 (42) | **0.034** | 0.099 | 0.133 | **<0.001** | 0.839 | 0.710 | 0.111 | 0.888 | 0.129 | 0.092 |
| *Head* | 13/133 (10) | 4/46 (9) | 4/37 (11) | 2/18 (11) | 0/20 (0) | 3/12 (25) | 0.746 | 0.766 | 0.174 | 0.123 | 0.973 | 0.127 | 0.222 | 0.126 | 0.317 | **0.019** |
| Thorax | 17/150 (11) | 2/48 (4) | 7/44 (16) | 3/21 (14) | 1/21 (5) | 4/16 (25) | 0.058 | 0.136 | 0.911 | **0.013** | 0.865 | 0.201 | 0.421 | 0.293 | 0.410 | 0.074 |
| *Axillary (vs. breast)* | 13/17 (76) | 1/2 (50) | 7/7 (100) | 3/3 (100) | 1/1 (100) | 1/4 (25) | **0.047** | 0.171 | 0.387 | 0.540 | - | - | **0.007** | - | **0.047** | 0.171 |
| Lymph node characteristics: size, $cm^2$ | 4 (2-9) | 4 (4-9) | 3 (1-4) | 5 (3-10) | 6 (4-29) | 4 (1-9) | **0.0288** | >0.9999 | >0.9999 | >0.9999 | 0.1069 | **0.002** | >0.9999 | >0.9999 | >0.9999 | 0.420 |
| Specimen appearance | | | | | | | | | | | | | | | | |
| Bloody (vs. chylous) | 123/150 (82) | 40/48 (83) | 39/44 (89) | 19/21 (90) | 14/21 (67) | 11/16 (69) | 0.466 | 0.438 | 0.122 | 0.209 | 0.823 | **0.033** | 0.068 | 0.060 | 0.095 | 0.893 |

Abbreviations: TB: tuberculosis; TBLs: tuberculous lymphadenitis; HIV: human immunodeficiency virus; ART: antiretroviral therapy; L: lymphotype; dTBLs: definite tuberculous lymphadenitis; nTBLs: non-tuberculous lyphadenitis;

**Table S5: Demographic, clinical, and microbiological differences between dTBL lymphotypes.** L3s had characteristics associated with more severe TBL. L3s were more likely to have HIV and larger lymph nodes compared to L1s and L2s. L2s were more likely to be female than L1s.

| **Characteristic**[¶] | **Total (n=89)** | **L1 (n=48) (*Prevotella-Corynebacterium*)** | **L2 (n=21) (Prevotella-*Streptococcus*)** | **L3 (n=20) (*Mycobacterium*)** | ***p-value* (L1 vs. L2)** | ***p-value* (L1 vs. L3)** | ***p-value* (L2 vs. L3)** |
|---|---|---|---|---|---|---|---|
| Age, years | 35 (29-40) | 33 (28-38) | 36 (28-46) | 37 (34-44) | >0.999 | 0.197 | 0.873 |
| Female | 48/89 (54) | 22/48 (46) | 15/21 (71) | 11/20 (55) | **0.050** | 0.491 | 0.275 |
| HIV | 49/89 (55) | 23/48 (48) | 11/21 (52) | 15/20 (75) | 0.733 | **0.040** | 0.133 |
| *CD4+* | 155 (76-251) | 157 (106-250) | 212 (64-385) | 92 (17-226) | >0.999 | 0.254 | 0.172 |
| *CD4+ <200 cells/µl* | 32/49 (65) | 16/23 (70) | 5/11 (45) | 11/15 (73) | 0.180 | 0.800 | 0.150 |
| *On ART* | 21/49 (43) | 11/23 (48) | 5/11 (45) | 5/15 (33) | 0.900 | 0.380 | 0.530 |
| Previous TB | 24/88 (27) | 11/47 (23) | 4/21 (19) | 9/20 (45) | 0.689 | 0.077 | 0.074 |
| Tobacco smoking | 21/89 (24) | 13/48 (27) | 4/21 (19) | 4/20 (20) | 0.480 | 0.540 | 0.940 |
| Antibiotic use within 1 year of recruitment | 22/87 (25) | 7/47 (15) | 6/20 (30) | 9/20 (45) | 0.153 | **0.008** | 0.327 |
| *At recruitment* | 10/22 (45) | 2/7 (29) | 2/6 (33) | 6/9 (67) | 0.850 | 0.130 | 0.200 |
| Lymph node characteristics: sites | | | | | | | |
| Neck | 78/89 (88) | 42/48 (88) | 17/21 (81) | 19/20 (95) | 0.480 | 0.350 | 0.170 |
| *Deep anterior cervical* | 36/78 (46) | 16/42 (38) | 7/17 (41) | 13/19 (68) | 0.826 | **0.028** | 0.101 |
| *Deep lateral cervical* | 15/78 (19) | 11/42 (26) | 2/17 (12) | 2/19 (11) | 0.230 | 0.170 | 0.910 |
| *Superficial* | 6/78 (8) | 5/42 (12) | 0/17 (0) | 1/19 (5) | 0.140 | 0.420 | 0.340 |
| *Supraclavicular* | 17/78 (22) | 7/42 (17) | 7/17 (41) | 3/19 (16) | **0.045** | 0.920 | 0.090 |
| *Head* | 4/78 (5) | 3/42 (7) | 1/17 (6) | 0/19 (0) | 0.860 | 0.230 | 0.280 |
| Thorax | 11/89 (12) | 6/48 (13) | 4/21 (19) | 1/20 (5) | 0.480 | 0.350 | 0.170 |
| *Axillary (vs. breast)* | 9/11 (82) | 6/6 (100) | 2/4 (50) | 1/1 (100) | 0.053 | - | 0.361 |
| Lymph node characteristics: size, $cm^2$ | 4 (2-9) | 4 (1-7) | 4 (1-4) | 8 (4-12) | 0.827 | **0.030** | **0.005** |
| Specimen appearance | | | | | | | |
| Bloody (vs. chylous) | 66/89 (74) | 38/48 (79) | 14/21 (67) | 14/20 (70) | 0.270 | 0.420 | 0.820 |

Abbreviations: TB: tuberculosis; TBLs: tuberculous lymphadenitis; HIV: human immunodeficiency virus; ART: antiretroviral therapy; L: lymphotype; dTBLs: definite tuberculous lymphadenitis.

**Figure S1: Paired analysis of controls and lymph fluid (n=33) indicates that environmental cross contamination is highly unlikely.** (A) α-diversity analyses show skin has higher diversity than lymph fluid. (B) β-diversity of lymph fluid differs to saline and skin. *DESeq2* volcano plots depicting differentially abundant taxa show that (C) lymph was enriched in *Mycobacterium* vs. (C) skin and (D) saline, and there were more differentially abundant taxa in skin vs. (C) lymph and (E) saline. Significantly more discriminatory taxa appear closer to the left or right, and higher above the threshold (red dotted line, FDR=0.2) as the degree of significance increases. Relative taxa abundance is indicated by circle size.

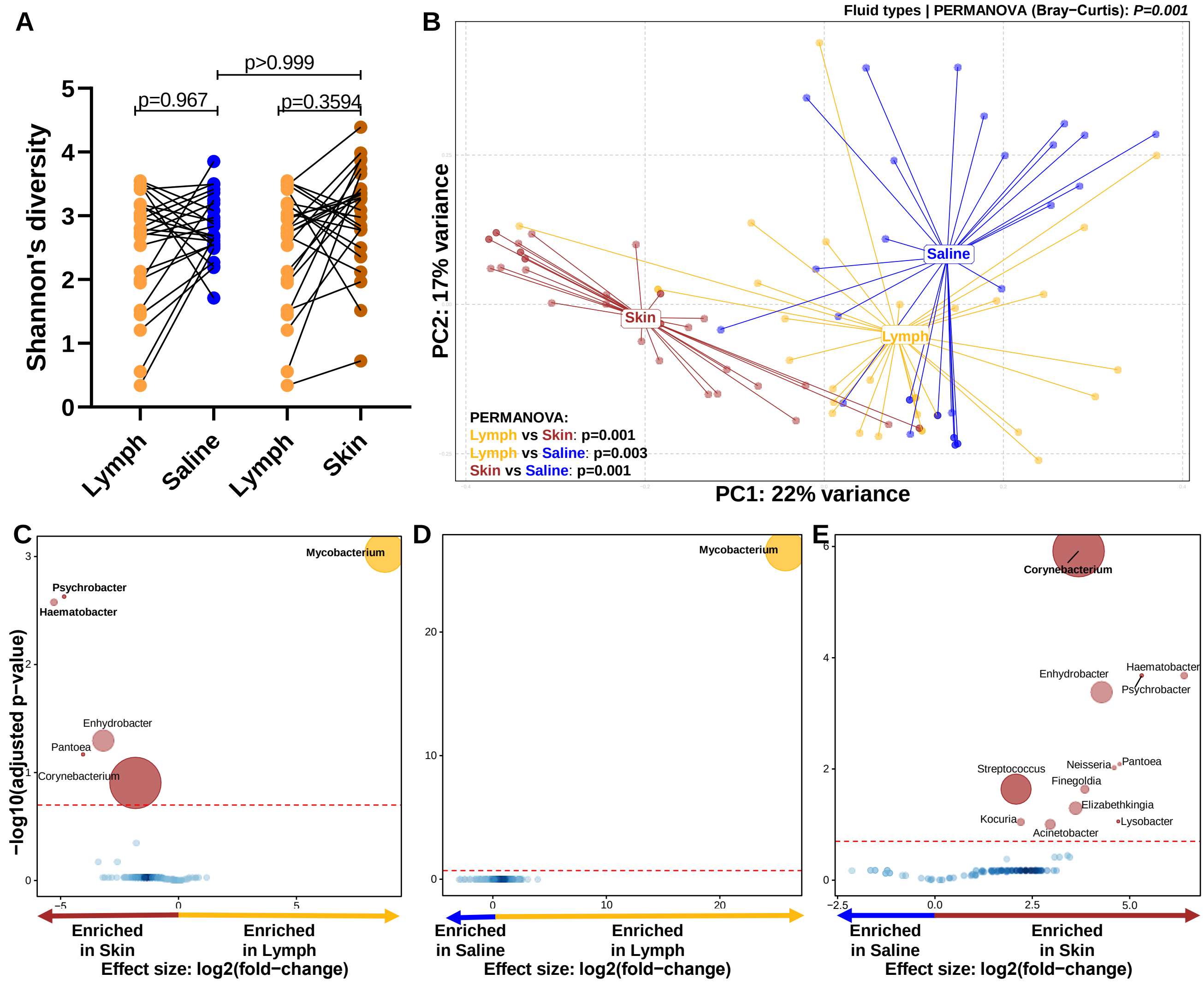

**Figure S2: *Mycobacterium* reads in FNABs of participants showing some nTBLs with Mycobacterium reads.** Relative abundance of *Mycobacterium* per participant stratified by TB status shows *Mycobacterium* in some nTBLs. Furthermore, not all dTBLs had detected *Mycobacterium* reads.

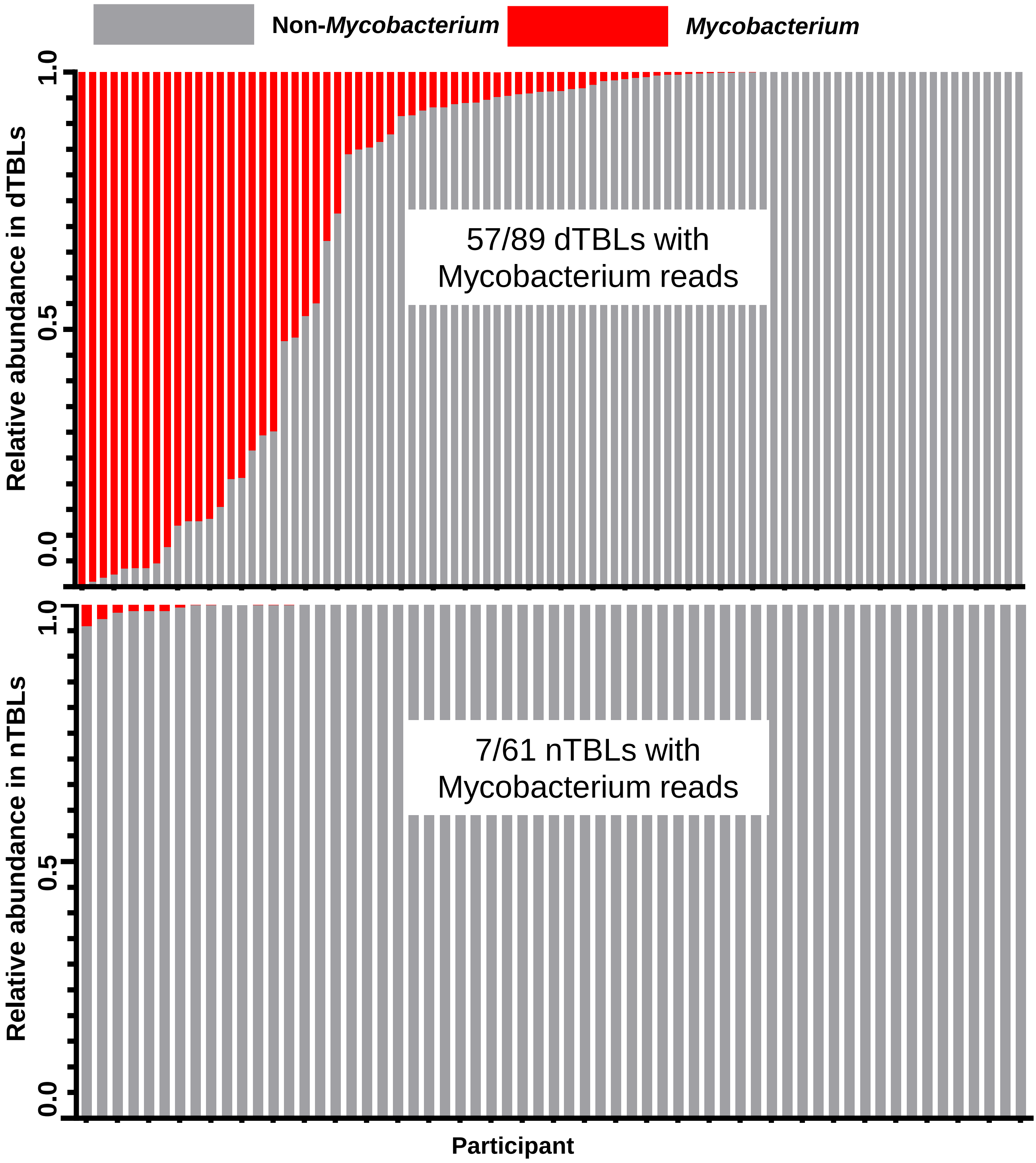

**Figure S3: Lymph node size is positively correlated to mycobacterial load.** In presumptive TBL patients, the size of the lymph node is associated with (A) relative abundance of mycobacterium genus reads present in the lymph node, and (B) Xpert and Ultra mycobacterial load. Xpert: Xpert MTB/RIF; Ultra: Xpert MTB/RIF Ultra; $r_s$: Spearman correlation coefficient.

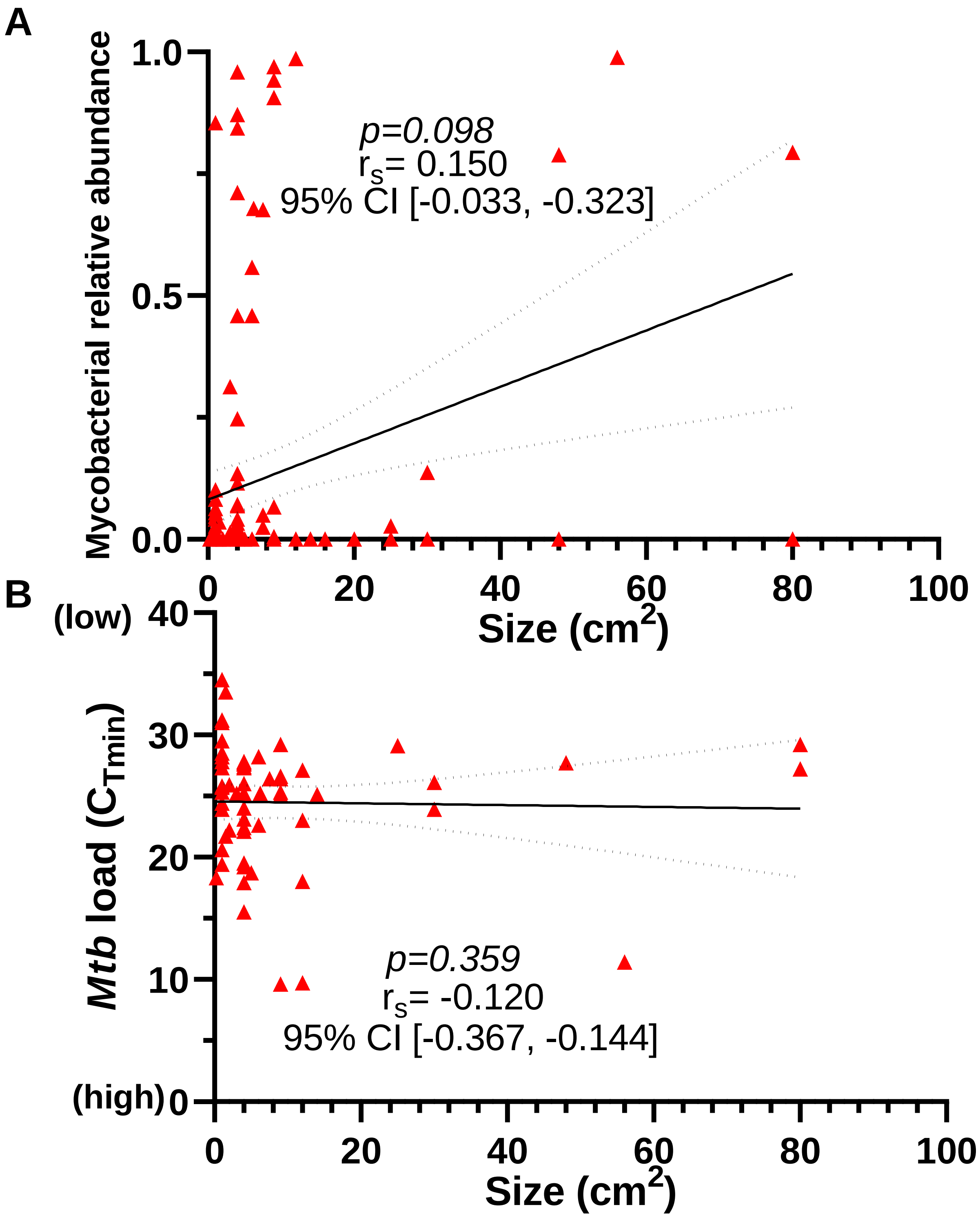

**Figure S4: HIV has a greater effect on the microbiome in patients co-infected with TB.** (A) Mycobacterial abundance did not differ by HIV status within dTBLs and within nTBLs, and (B) HIV-positive dTBLs were enriched in *Mycobacterium* compared to nTBLs. Circle sizes represent relative abundances. dTBLs: definite tuberculous lymphadenitis; nTBLs: non-tuberculous lymphadenitis.

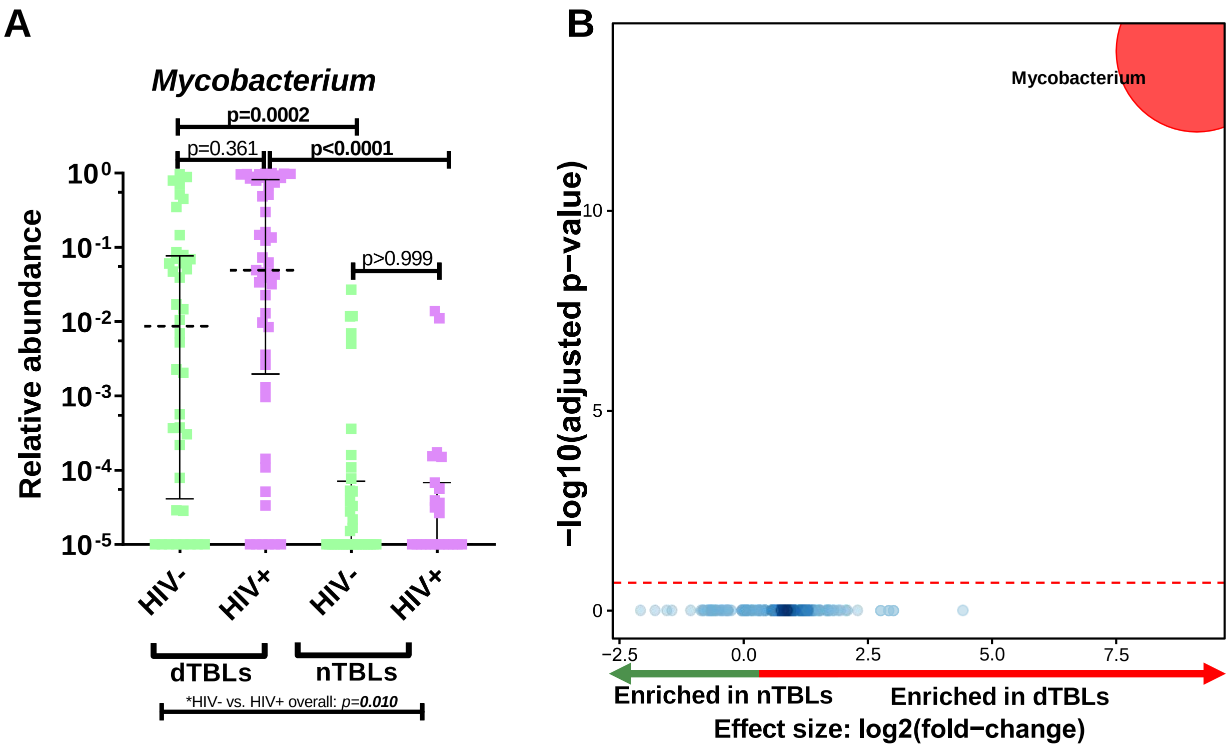

**Figure S5: Five microbial community states observed in presumptive TBL patients are enriched with distinct taxa.** L1 had no enriched taxa, and was depleted in (A) *Enhydrobacter*, (B) *Mycobacterium*, and (C) *Streptococcus*, *Anaerosinus*, *Neisseria* and *Kocuria*. L3 was enriched in (D) *Acinetobacter* and depleted of *Prevotella*. L5 was enriched in *Streptococcus* accompanied with (E) *Anaerosinus*, *Neisseria*, *Kocuria* and *Prevotella* vs. L2, and with (F) *Bacteroides* and *Kocuria* vs. L3. Significantly more discriminatory taxa (bolded) appear closer to the left or right, and higher above the threshold (red dotted line, FDR=0·2) as significance increases. Relative abundance of taxa is indicated by circle size. L: Lymphotype.

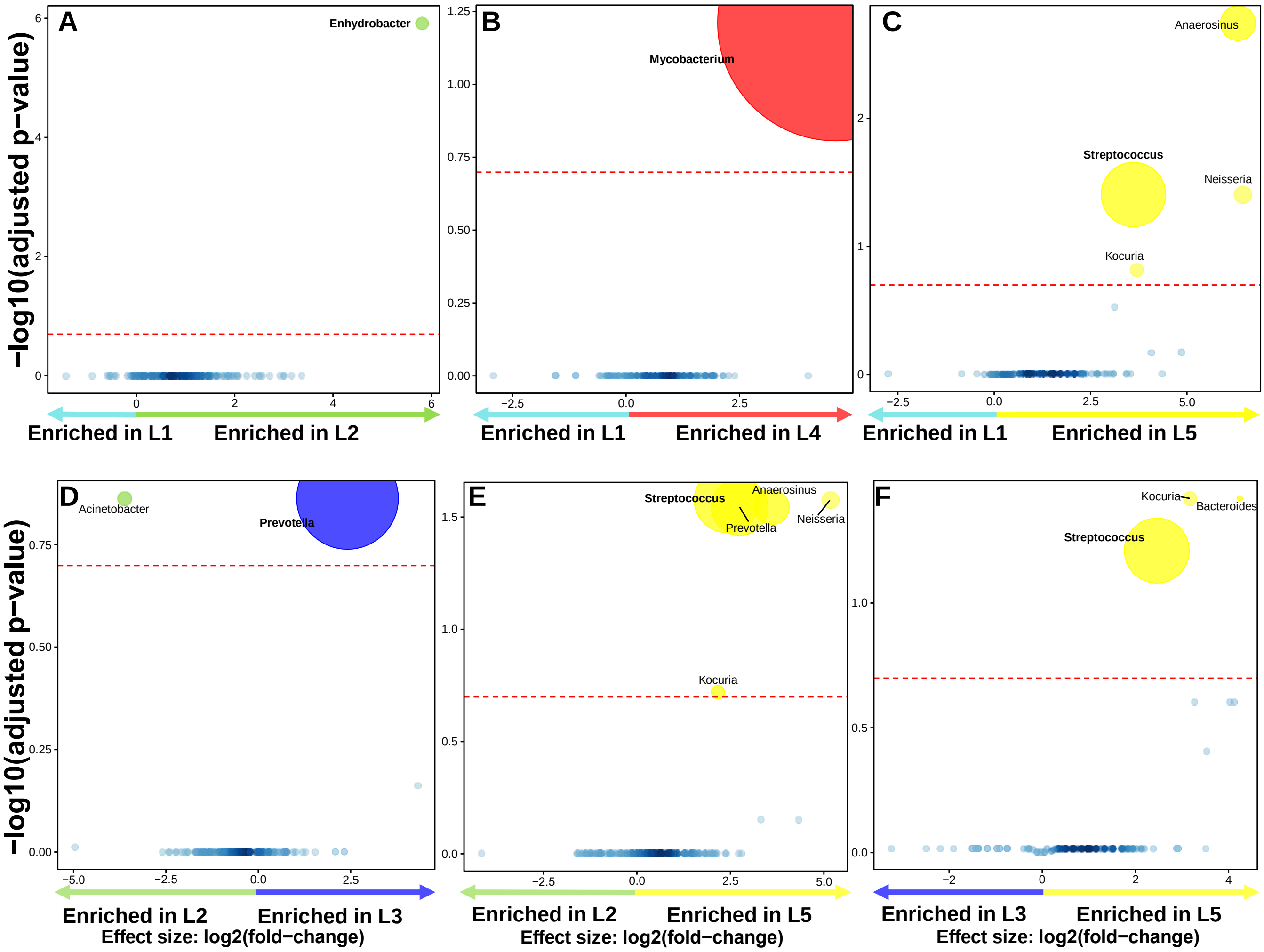

**Figure S6: The Laplace approximation of model evidence is a measure of the model fit**. Laplace approximation predicts no clustering for nTBL patients. Lower values indicate better fit. nTBLs: non-tuberculous lymphadenitis.

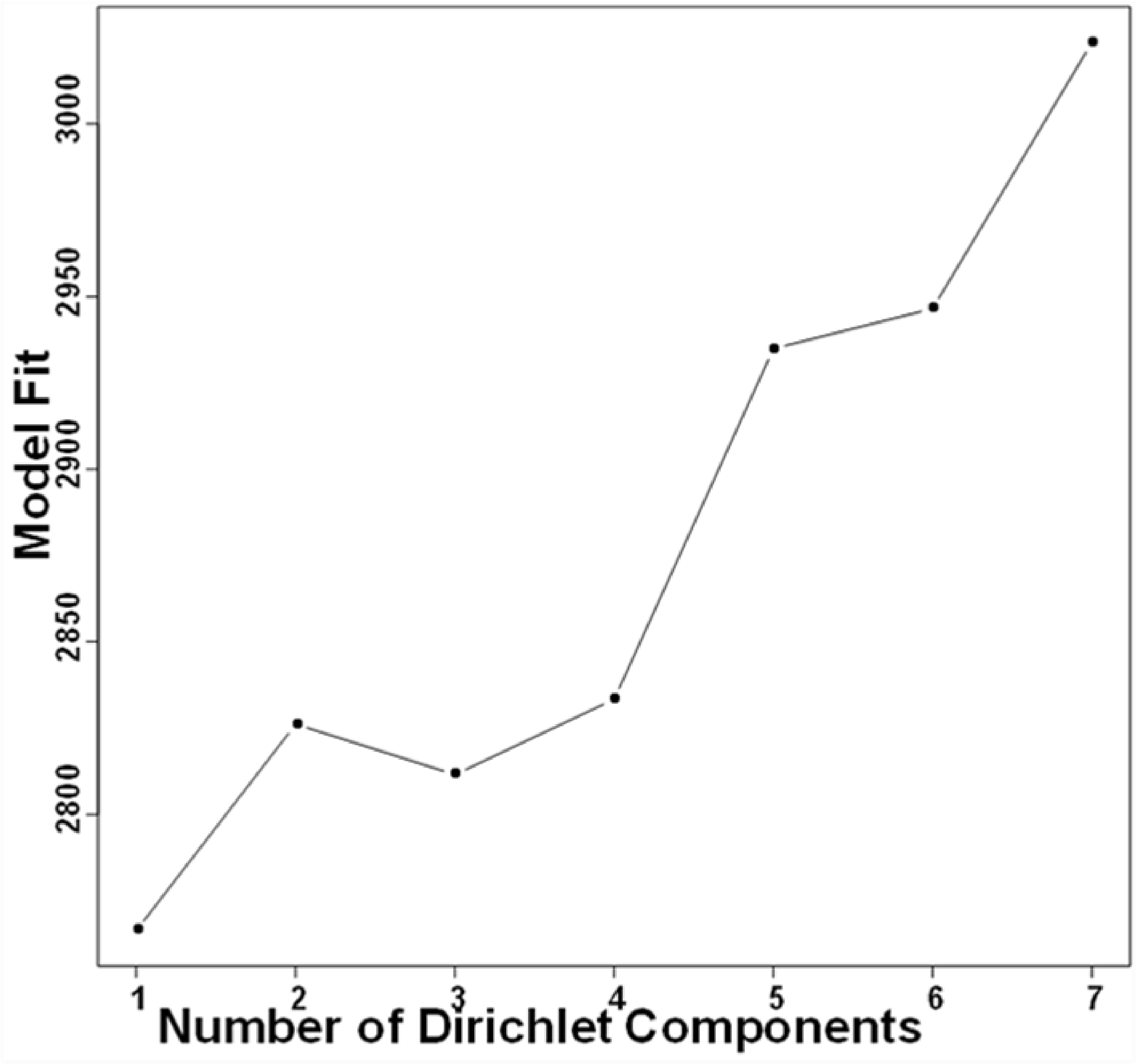

**Figure S7: Predicted metagenome function in HIV-positive nTBLs versus HIV-negative nTBLs.** Volcano plot depicting functional pathways differing between HIV-positive and HIV-negative nTBLs. Significantly more discriminatory pathways appear closer to the left or right, and higher above the threshold (red dotted line, FDR=0.05). Key pathways of interest include "cell cycle - *Caulabacter*", "bacterial secretion system", "taurine and hypotaurine metabolism", and "histidine metabolism". Relative gene abundance is indicated by circle size.

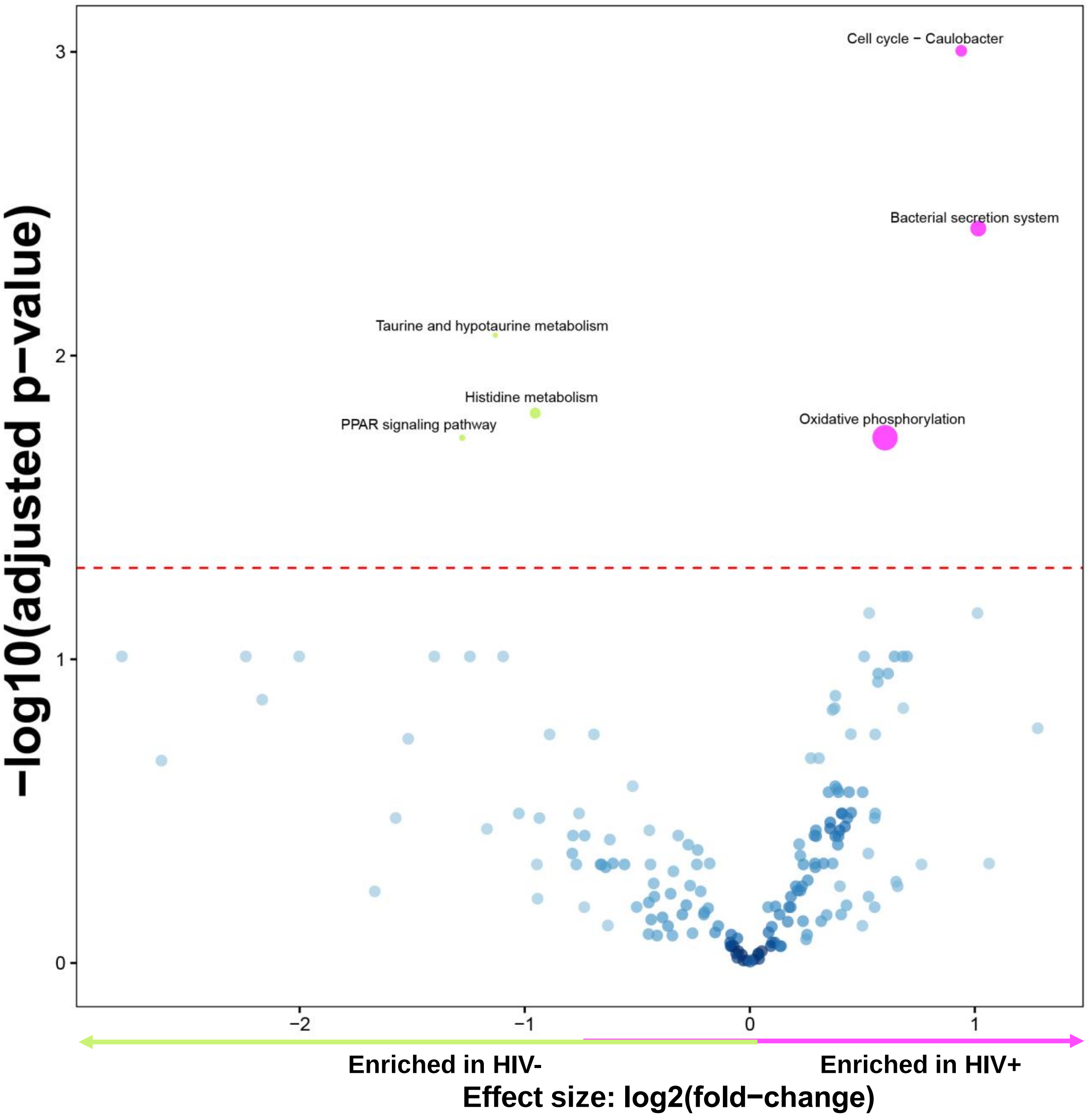

**Figure S8: Inferred metagenomes of lymphotypes in all patients.** Volcano plot depicting differentially enriched pathways in L4 included pathways involving lipid biosynthesis, fatty acids, and SCFA metabolism i.e. lipid biosynthesis proteins, propanoate metabolism, benzoate degradation, and valine, leucine and isoleucine degradation. Significantly more discriminatory pathways appear closer to the left or right, and higher above the threshold (red dotted line, FDR=0.05). Relative gene abundance is indicated by circle size. L: Lymphotype.

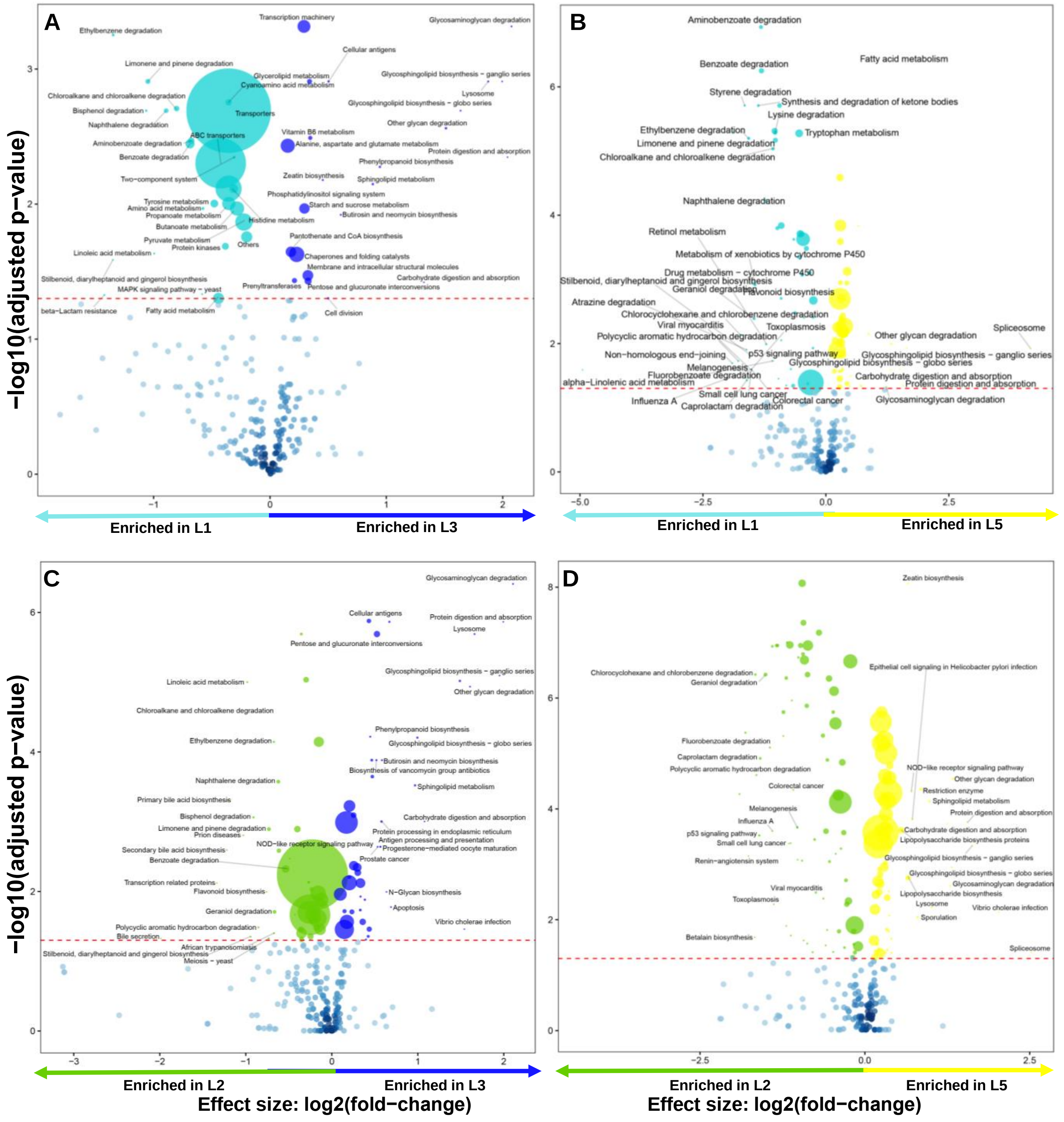

**Figure S8 cont.**

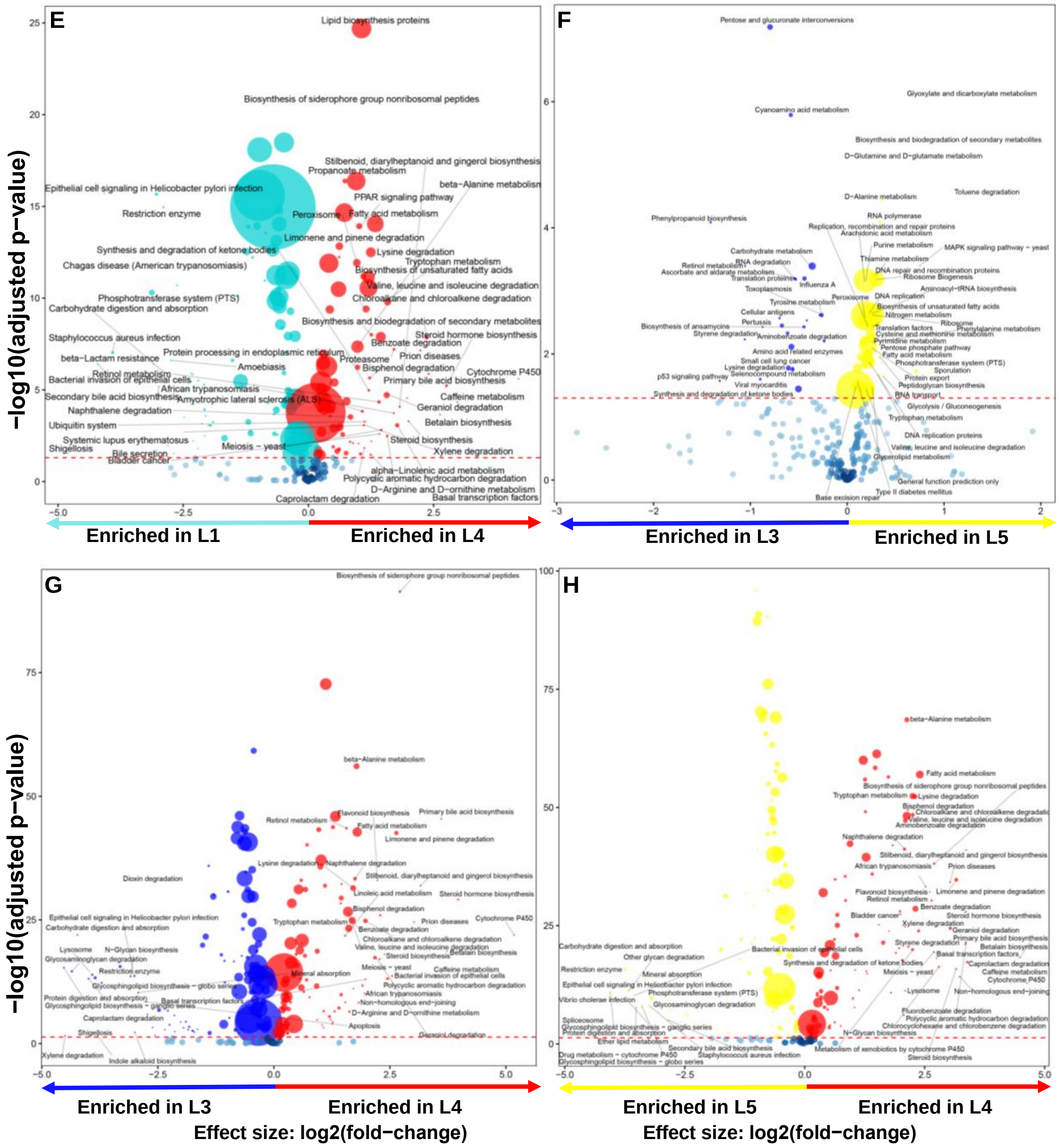

**Figure S9: Inferred metagenomes of lymphotypes in dTBLs.** Volcano plot depicting differentially enriched pathways in L3 included pathways involving lipid biosynthesis, fatty acids, and SCFA metabolism i.e. lipid biosynthesis proteins, propanoate metabolism, benzoate degradation, and valine, leucine and isoleucine degradation. Significantly more discriminatory pathways appear closer to the left or right, and higher above the threshold (red dotted line, FDR=0.05). Relative gene abundance is indicated by circle size. L: Lymphotype.

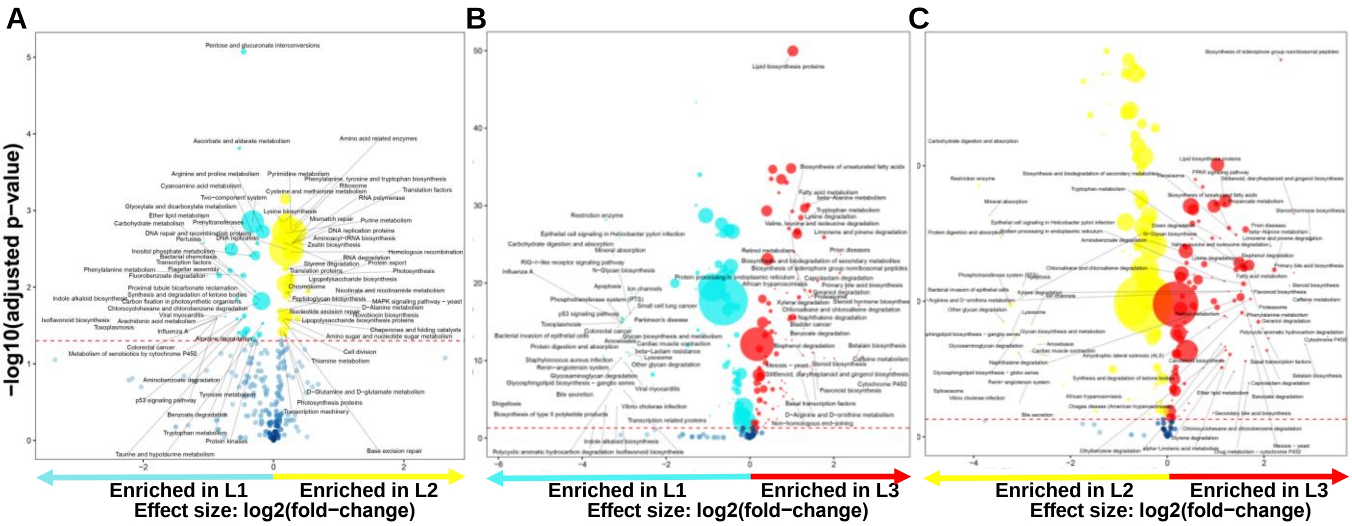
